# A new perspective on the evolution of the interaction between the Vg/VGLL1-3 proteins and the TEAD transcription factors

**DOI:** 10.1101/2020.05.21.107789

**Authors:** Yannick Mesrouze, Gustavo Aguilar, Fedir Bokhovchuk, Typhaine Martin, Clara Delaunay, Frédéric Villard, Marco Meyerhofer, Catherine Zimmermann, Patrizia Fontana, Roman Wille, Thomas Vorherr, Dirk Erdmann, Pascal Furet, Clemens Scheufler, Tobias Schmelzle, Markus Affolter, Patrick Chène

## Abstract

The most downstream elements of the Hippo pathway, the TEAD transcription factors, are regulated by several cofactors, such as Vg/VGLL1-3. Earlier findings on human VGLL1 and here on human VGLL3 show that these proteins interact with TEAD via a conserved amino acid motif called the TONDU domain. Surprisingly, our studies reveal that the TEAD-binding domain of *Drosophila* Vg and of human VGLL2 is more complex and contains an additional structural element, an Ω-loop, that contributes to TEAD binding and *in vivo* function. To explain this unexpected structural difference between proteins from the same family, we propose that, after the genome-wide duplications at the origin of vertebrates, the Ω-loop present in an ancestral *VGLL* gene has been lost in some VGLL variants. These findings illustrate how structural and functional constraints can guide the evolution of transcriptional cofactors to preserve their ability to compete with other cofactors for binding to transcription factors.

## Introduction

The Hippo pathway, which controls organ size and tissue regeneration via the regulation of cell division and apoptosis (Ma, Meng et al., 2019, Wang, Yu et al., 2017), is an intense field of research in regenerative medicine (Fu, Plouffe et al., 2017, Hong, Meng et al., 2016, Moya & Halder, 2018, Wang, Liu et al., 2018). Furthermore, deregulation of Hippo signalling in several cancers suggests that molecules targeting this pathway may have antineoplastic properties (Janse van Rensburg & Yang, 2016, Maugeri-Sacca & De Maria, 2018, Park, Shin et al., 2018, Santucci, Vignudelli et al., 2015, Zanconato, Cordenonsi et al., 2016) (but see (Kakiuchi-Kiyota, Schutten et al., 2019, Moya, Castaldo et al., 2019)). The Hippo pathway regulates, via a signalling cascade involving the MST1/2 and LATS1/2 protein kinases, the TEAD (TEA/ATTS domain) transcription factors that, upon binding to YAP (Yes associated protein) or VGLL1-3 (Vestigial-like), become transcriptionally active (Landin-Malt, Benhaddou et al., 2016, Lin, Park et al., 2017, Liu, Wong et al., 2012, Ma et al., 2019). The elucidation of the structures of the YAP:TEAD (Chen, Chan et al., 2010, Li, Zhao et al., 2010) and of the VGLL1:TEAD (Pobbati, Chan et al., 2012) complexes has shed some light on how these proteins bind to TEAD. The TEAD-binding domain of YAP is formed of a β-strand:α-helix:Ω-loop motif while the TEAD-binding domain of VGLL1 contains only a β-strand:α-helix motif, known as TONDU domain, which binds to TEAD in a fashion similar to the β-strand:α-helix region of YAP (Fig S1) (Li et al., 2010). The residues present in the TONDU domain of VGLL1 are well-conserved amongst the members of the Vg/VGLL1-3 family (Vg, vestigial, *Drosophila* homolog of VGLL1-3) (Simon, Faucheux et al., 2016) and no other regions of these proteins outside this motif have so far been reported to play a direct role in the interaction with TEAD proteins. Therefore, the main structural difference between the TEAD-binding domain of these two cofactor families is the presence of an Ω-loop in the Yki/YAP proteins (Yki, Yorkie, *Drosophila* homolog of YAP), but not in the Vg/VGLL1-3 proteins. However, we made an observation which suggests that some members of the Vg/VGLL1-3 family may also possess an Ω-loop in their TEAD-binding domain. We identified a region in the amino acid sequence of the Vg protein from *Drosophila melanogaster*, residues 323-334 (Vg^323-334^), which has a strong homology with the Ω-loop of YAP and is separated from the TONDU domain (Vg^288-311^) by a similar number of residues compared with the Ω-loop from the β-strand:α-helix region in YAP (Fig 1A). This unexpected finding, which suggests that Vg contains an Ω-loop, echoes earlier findings. Simmonds et al. have mapped the Sd-binding domain of Vg (Sd, Scalloped, *Drosophila* homolog of human TEAD) to Vg^279-335^, which contains not only the TONDU domain but also the putative Ω-loop, Vg^323-334^ (Simmonds, Liu et al., 1998). However, these authors did not conduct experiments with shorter fragments to determine whether the entire Vg^279-335^ is required for efficient binding to Sd. In their study of the ladybird beetle Vg protein, Ohde et al. observed that, in addition to the TONDU domain, this protein also contains a small sequence conserved in the Vg protein of other species, including *D. melanogaster* (Ohde, Masumoto et al., 2009). This sequence corresponds to the putative Ω-loop, Vg^321-334^, but these authors report that the function of this region is unknown. Altogether, these observations prompted us to investigate the interaction between Vg and Sd in more detail and, in the light of our findings, to revisit the interaction between TEAD and VGLL1-3.

**Figure 1.**
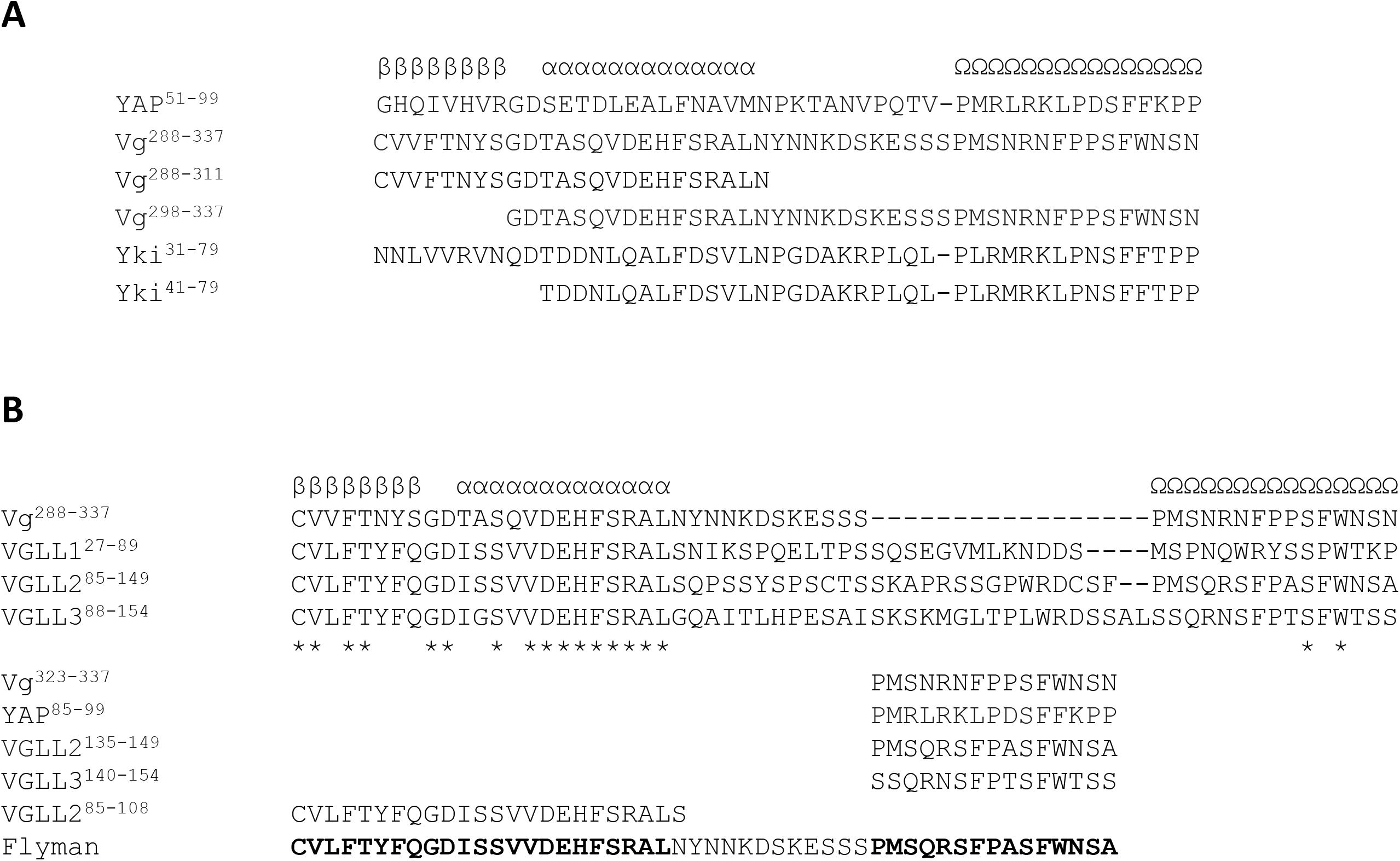
Primary sequence alignments. **A.** The structures of the YAP:TEAD (PDB 3KYS (Li et al., 2010)) and of the VGLL1:TEAD (PDB 5Z2Q (Pobbati et al., 2012)) complexes were used to manually align the primary sequences of YAP (UniProt P46937), Vg (UniProt Q26366) and Yki (UniProt Q45VV3). The secondary structure adopted by YAP upon binding to TEAD is indicated. β: β-strand; α: α-helix; Ω: Ω-loop. Gaps are indicated by dashes (-). **B.** The primary sequences of human VGLL1 (UniProt Q99990), VGLL2 (UniProt Q8N8G2) and VGLL3 (UniProt A8MV65) were manually aligned to the primary sequence of Vg to obtain an optimal alignment of the residues in the β-strand:α-helix and of the conserved serine and tryptophan residues located near the C-terminus. The asterisks (*) indicate strictly conserved residues. The primary sequence of the peptides used in the TR-FRET assay are given at the bottom of the figure. The residues in bold and normal fonts in the sequence of Flyman are from VGLL2 and Vg, respectively.

## Results

### Biophysical characterization of the Vg:Sd interaction

The alignment of the primary sequence of TEAD and Sd and the analysis of the structure of the YAP/VGLL1:TEAD complexes (Chen et al., 2010, Li et al., 2010, Pobbati et al., 2012) allowed us to tentatively map the Vg/Yki-binding domain of Sd between residues 223 and 440 (Fig S2). This protein fragment, hereafter referred to as wt^Sd^, was purified from *Escherichia coli* as a mixture of acylated and non-acylated proteins (Fig S3). The acylation of the human TEAD proteins has been described (Chan, Han et al., 2016, Mesrouze, Meyerhofer et al., 2017b, Noland, Gierke et al., 2016) and the physiological role of Sd acylation on tissue overgrowth has been shown in *Drosophila* (Chan et al., 2016). A circular dichroism (CD) analysis shows that recombinant wt^Sd^ is well-folded and its CD spectrum is similar to that of TEAD4^217-434^ (corresponding TEAD4 fragment) (Fig S4A). The structure of the YAP:TEAD complex (Li et al., 2010) and the alignment of the amino acid sequence of YAP and Yki enabled us to delineate the Sd-binding domain of Yki to Yki^31-79^ (Fig 1A). The affinities (K_d_) of Vg^288-311^ (β-strand:α-helix, TONDU domain) and Yki^31-79^ (β-strand:α-helix:Ω-loop) for wt^Sd^, as measured by Surface Plasmon Resonance (SPR), are 291 and 23 nM, respectively (Table 1). These K_d_ values are similar to those of their counterparts VGLL1^27-51^ (160 nM) and YAP^50-99^ (11 nM) measured with TEAD4 (Hau, Erdmann et al., 2013, Mesrouze, Hau et al., 2014).

**Table 1.**
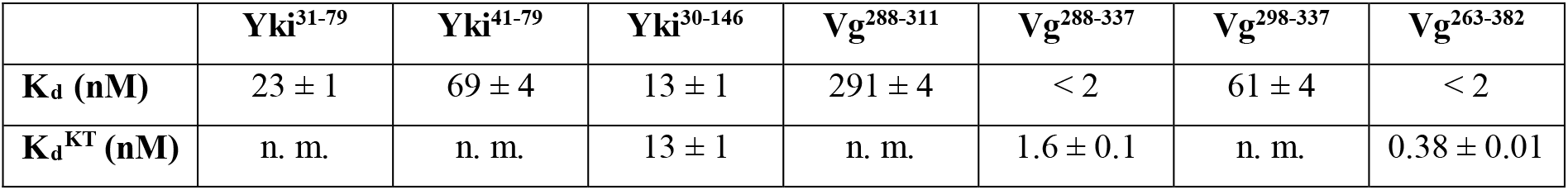
Affinity of different Yki and Vg fragments for wt^Sd^. N-biotinylated-Avitagged wt^Sd^ was immobilized on sensor chips and the affinity of different Yki and Vg fragments was measured by Surface Plasmon Resonance (SPR). Affinities were determined either at equilibrium by a standard SPR method (K_d_) or by kinetic titration (K_d_^KT^). The average and standard error of n ≥ 2 separate experiments are given. n. m.: not measured.

|  | Yki<sup>31-79</sup> | Yki<sup>41-79</sup> | Yki<sup>30-146</sup> | Vg<sup>288-311</sup> | Vg<sup>288-337</sup> | Vg<sup>298-337</sup> | Vg<sup>263-382</sup> |
| --- | --- | --- | --- | --- | --- | --- | --- |
| **K<sub>d</sub> (nM)** | 23 ± 1 | 69 ± 4 | 13 ± 1 | 291 ± 4 | < 2 | 61 ± 4 | < 2 |
| **K<sub>d</sub><sup>KT</sup> (nM)** | n. m. | n. m. | 13 ± 1 | n. m. | 1.6 ± 0.1 | n. m. | 0.38 ± 0.01 |

We next studied Vg^288-337^, which contains the TONDU domain and the putative Ω-loop (Vg^323-334^). Binding of Vg^288-337^ to wt^Sd^ was so tight (K_d_ < 2 nM) that we were unable to accurately determine its affinity using a standard SPR procedure. Therefore, we used the kinetic titration method, which allows K_d_ determination for high affinity ligands (Karlsson, Katsamba et al., 2006). To validate this methodology for our proteins, we purified Yki^30-146^, which according to results obtained with the equivalent YAP^50-171^ fragment should have a high affinity for wt^Sd^ (Hau et al., 2013). Since the K_d_ values of Yki^30-146^ measured by both SPR methods are similar (Table 1), kinetic titration was applied to Vg^288-337^ and a K_d_^KT^ = 1.6 nM was determined (Table 1). Vg^288-337^ binds 190 times more tightly to wt^Sd^ than the TONDU domain alone (Vg^288-311^), showing that the region Vg^312-337^ (including the putative Ω-loop) dramatically enhances the affinity for wt^Sd^. To determine whether additional regions outside Vg^288-337^ contribute to the binding to wt^Sd^, the longer protein fragment Vg^263-382^ was purified. The affinity of Vg^263-382^ – K_d_^KT^ = 0.38 nM (Table 1) – is similar (fourfold difference) to that of Vg^288-337^, suggesting that no other regions from Vg^263-382^ contribute significantly to the interaction with wt^Sd^. The residues in the β-strand of the Sd/TEAD-binding domain of YAP/Yki are variable across species and their contribution to the interaction with Sd/TEAD is limited (Hau et al., 2013, Li et al., 2010). In contrast, the amino acids from the β-strand in the Sd/TEAD-binding domain of Vg/VGLL1-3 are well conserved (Simon et al., 2016) suggesting that they might contribute more to the interaction with these transcription factors. The K_d_ values for wt^Sd^ of Yki^41-79^ and Vg^298-337^, which both lack the putative β-strand region (Fig 1A), are 69 nM and 61 nM, respectively (Table 1). As observed for YAP, the deletion of the putative β-strand of the Sd-binding domain of Yki has little effect on binding to wt^Sd^ (only a threefold change in Kd). In contrast, the deletion of the putative β-strand from the Sd-binding domain of Vg leads to a 40-fold reduction in binding, indicating that this secondary structure element plays an important part in the interaction with wt^Sd^.

### Structural characterization of the Vg:Sd interaction

To study the interaction between Vg and Sd at the atomic level, we set up co-crystallization experiments between wt^Sd^ and Vg^288-337^ as well as Vg^298-337^ and we obtained crystals of the Vg^298-337^:wt^Sd^ complex diffracting at 1.85 Å (PDB 6Y20; Table S1). In agreement with their high amino acid sequence homology and in line with our CD data (Fig S4A), Sd and TEAD have a very similar three-dimensional structure (Fig S6). The superimposition of the structure of the Vg:Sd complex on that of the YAP:TEAD and VGLL1:TEAD complexes show that the α-helix of the Sd/TEAD-binding site of these three co-factors bind to a similar location at the surface of Sd/TEAD (Fig 2A). The axes of the α-helix of Vg and YAP are not aligned, as also previously noticed between VGLL1 and YAP (Mesrouze, Erdmann et al., 2016), but the α-helices of Vg and VGLL1 bind in a similar fashion (Fig 2B). The three conserved residues in the VxxHF motif from the TONDU domain of VGLL1 and Vg occupy the same position and His305^Vg^ forms the same two hydrogen bonds with Sd (with Ser342^Sd^ and Val395^Sd^) as His44^VGLL1^ does with TEAD in the VGLL1:TEAD complex (Pobbati et al., 2012) (Fig 2C). The structure of the Vg:Sd complex shows that the Sd-binding domain of Vg indeed contains an Ω-loop, which binds to Sd in a similar fashion than the Ω-loop of YAP binding to TEAD (Fig 2A). The linker region connecting the α-helix and the Ω-loop of Vg is not completely resolved in our structure probably because of its flexibility (Fig 2A). The three hydrophobic residues Met324^Vg^:Phe329^Vg^:Phe333^Vg^ present in the Ω-loop of Vg occupy the same position at the binding interface as Met86^YAP^:Leu91^YAP^:Phe95^YAP^ do (Fig 2D). They also create a hydrophobic core within the bound Ω-loop, which is stabilized in the YAP:TEAD complex by Phe96^YAP^ that makes a π-cation interaction with Arg87^YAP^ (Fig 2E). In the Vg:Sd complex, Trp334^Vg^ is at the position of Phe96^YAP^ but it does not engage in a π-cation interaction because Ser325^Vg^ replaces Arg87^YAP^. This suggests that the presence of a larger aromatic residue at this position might be sufficient to stabilize and shield the hydrophobic core of the bound Ω-loop from solvent. The key salt bridge between Arg89^YAP^ and Asp272^TEAD4^ found in the YAP:TEAD complex (Mesrouze, Bokhovchuk et al., 2017a) is also observed between Arg327^Vg^ and Asp276^Sd^ in the Vg:Sd complex (Fig 2F). Finally, Ser332^Vg^ is within hydrogen bond distance of Glu267^Sd^:Tyr435^Sd^, as is the case for Ser94^YAP^ and Glu263^TEAD4^:Tyr409^TEAD4^ in the YAP:TEAD complex (Fig 2F). Overall, the structure of the Vg^298-337^:Sd^223-440^ complex clearly demonstrates the presence of an Ω-loop in the Sd-binding domain of Vg and that it interacts with Sd in a manner similar to that of the Ω-loop of YAP with TEAD.

**Figure 2.**
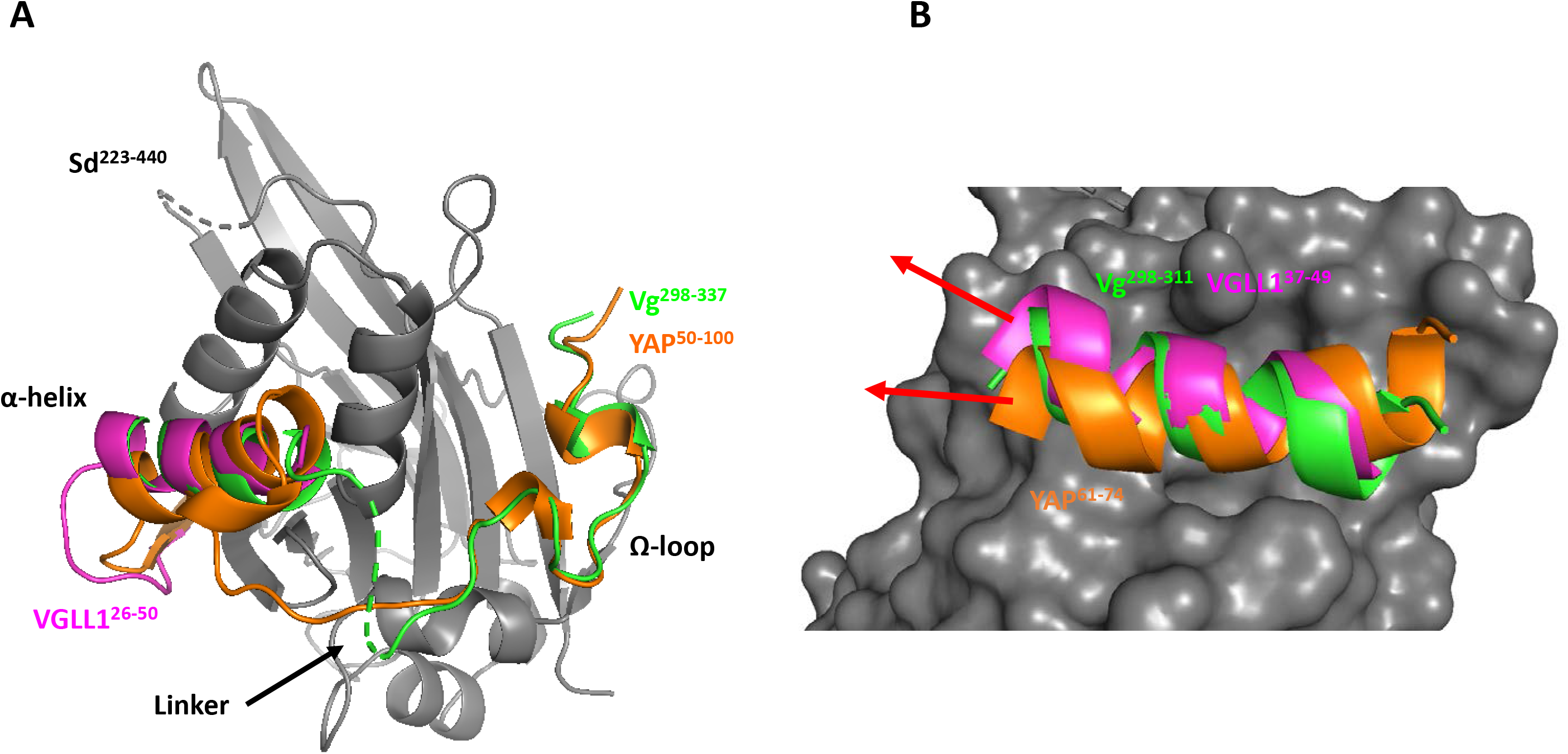
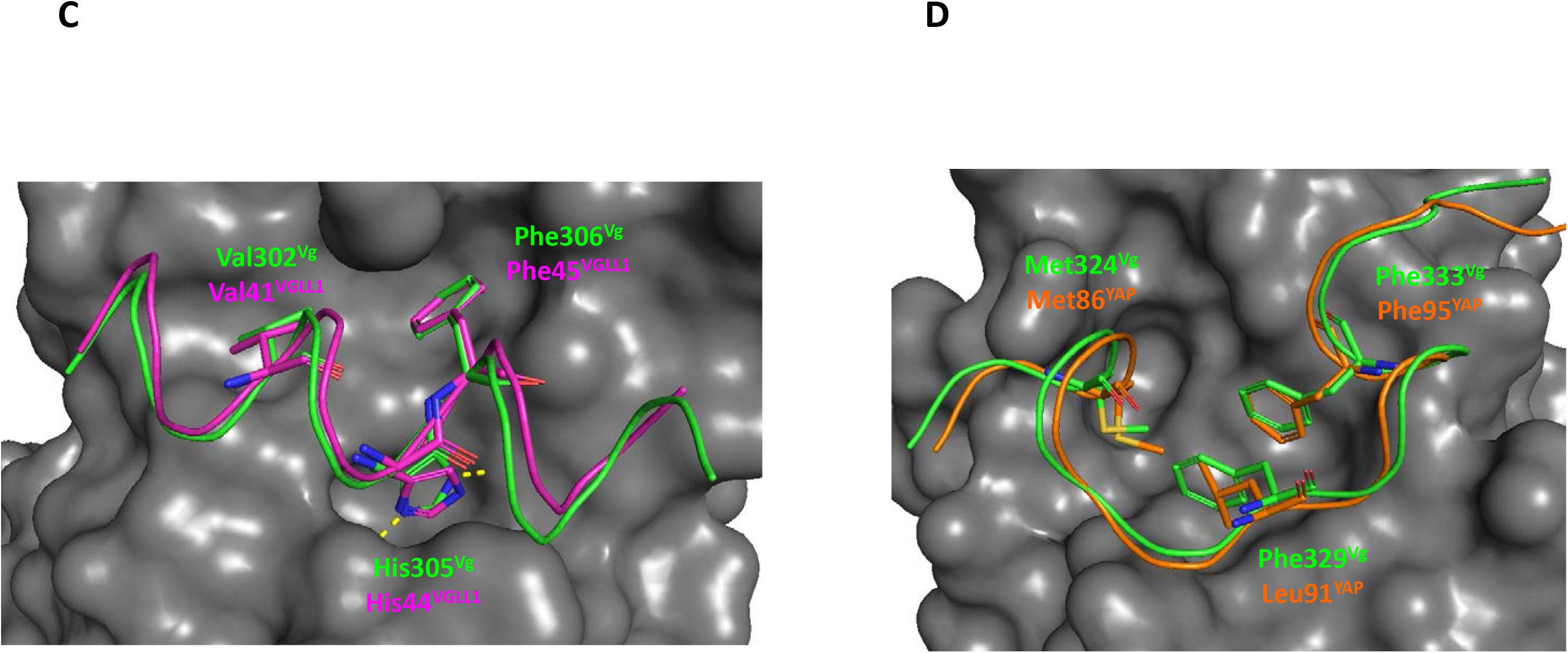
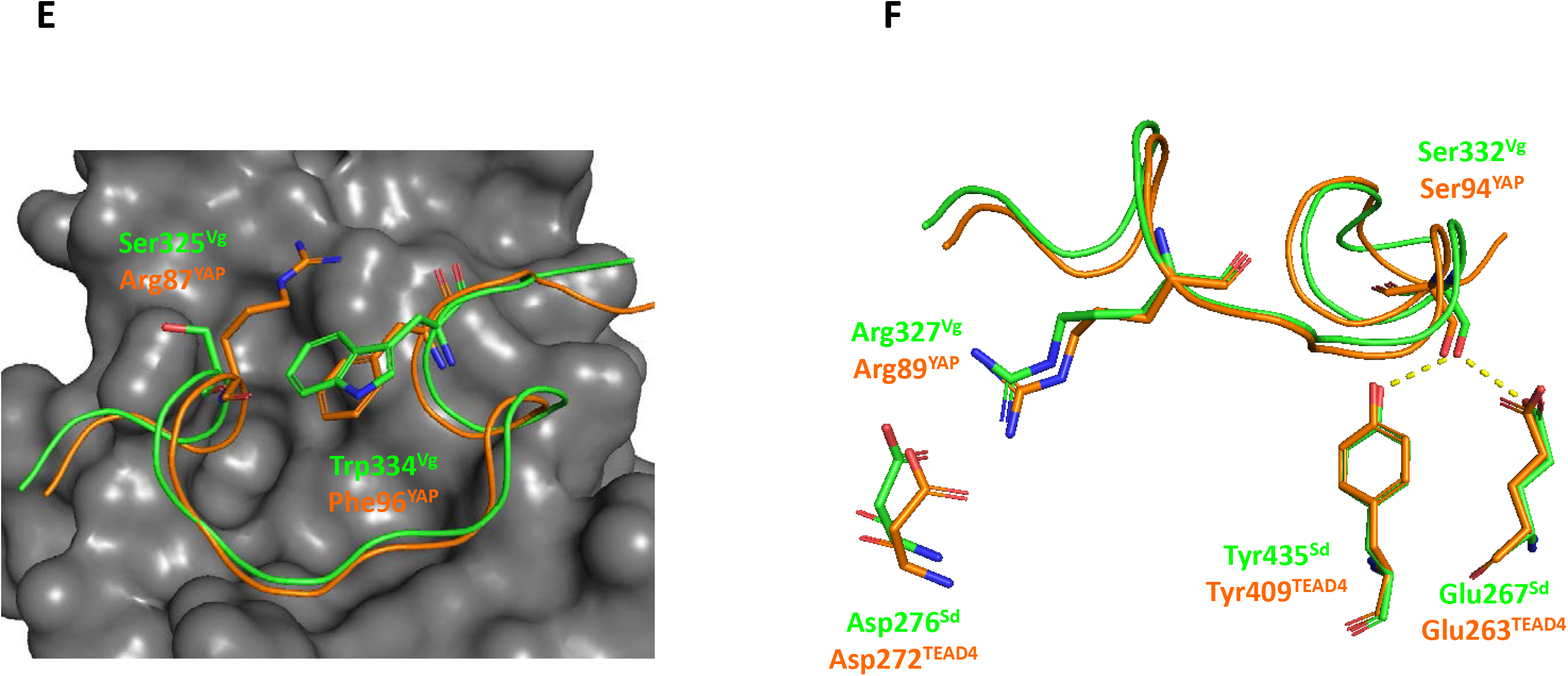
Structure of the Vg:Sd complex. The structure of the Vg^298-337^:wt^Sd^ complex (PDB 6Y20) is superimposed on that of the YAP:TEAD (PDB 3KYS (Li et al., 2010)) and VGLL1:TEAD (PDB 5Z2Q (Pobbati et al., 2012)) complexes. **A.** wt^Sd^, Vg^298-337^, VGLL1^26-50^ and YAP^50-100^ are represented in grey, green, magenta and orange, respectively. **B, C**. α-helix binding pocket. The α-helices present in the Sd/TEAD-binding domain of Vg, VGLL1 and YAP are represented in green, magenta and orange, respectively. Sd is in grey. The red arrows indicate the approximate axes of the α-helices. The three conserved residues of the VxxHF motif are represented by sticks and the hydrogen bonds made between His305^Vg^ and Sd are indicated by yellow dots. **D-F**. Ω-loop binding pocket. The Ω-loop present in the Sd/TEAD-binding domain of Vg and YAP are represented in green and orange, respectively. Sd is in grey. The different interactions are described in the text. The residue labelled Met324^Vg^ is a norleucine. This amino acid was introduced in Vg^298-337^ to enhance its stability against oxidation. The yellow dots indicate hydrogen bonds. The figures were drawn with PyMOL (Schrödinger Inc., Cambridge, MA).

### Effect of mutations in the Ω-loop binding pocket of Sd

The β-strand:α-helix region of YAP (YAP^44-74^) has a low affinity for TEAD (Mesrouze et al., 2014) while its Ω-loop (YAP^85-99^), binds more tightly to it (Hau et al., 2013, Zhang, Lin et al., 2014). Consequently, mutations in the Ω-loop binding pocket of TEAD are more destabilizing than mutations in the β-strand:α-helix binding pocket (Li et al., 2010, Mesrouze et al., 2017a). Since the β-strand:α-helix region of Vg (Vg^288-311^) has a high affinity for wt^Sd^ (Table1), we hypothesized that the Vg:Sd complex might be more permissive to mutations in the Ω-loop binding pocket of Sd than the Yki:Sd complex, which should behave like the YAP:TEAD complex. To test this hypothesis, we engineered mutations disrupting key interactions at the Ω-loop binding interface. The Asp276Ala^Sd^ mutation prevents the formation of a salt bridge with Arg327^Vg^ in the Vg:Sd complex (Fig 2F) and probably with Arg69^Yki^ in the Yki:Sd complex. The same mutation in TEAD4, Asp272Ala^TEAD4^, has a strong destabilizing effect on the YAP:TEAD complex (Mesrouze, Bokhovchuk et al., 2018). The Tyr435His^Sd^ deletes the hydrogen bond with Ser332^Vg^ in the Vg:Sd complex (Fig 2F) and probably with Ser74^Yki^ in the Yki:Sd complex. This mutation in TEAD1, Tyr421His^TEAD1^, which is at the root of Sveinsson’s chorioretinal atrophy (Fossdal, Jonasson et al., 2004), has a major negative impact on the YAP:TEAD complex (Bokhovchuk, Mesrouze et al., 2019). Asp246Ala^Sd^ and Tyr435His^Sd^ were purified, and their CD spectrum shows that the mutations do not affect the structure of Sd (Fig S4B) as previously observed with the corresponding TEAD4 mutations (Bokhovchuk et al., 2019, Mesrouze et al., 2017a). The affinity of Vg^263-382^ and of Yki^30-146^ for Asp276Ala^Sd^ is reduced by 1.58 and 3.6 kcal/mol, respectively (Table 2) and the Tyr435His^Sd^ mutation lowers the binding of Vg^263-382^ and of Yki^30-146^ by 2.44 and 4.11 kcal/mol, respectively (Table 2). Therefore, the Vg:Sd complex is less destabilized by these two mutations than the Yki:Sd complex, suggesting that the higher affinity of the β-strand:α-helix region of Vg for wt^Sd^ mitigates the impact of disruptive mutations at the Ω-loop binding pocket.

**Table 2.** Effect of the Asp276Ala^Sd^ and Tyr435His^Sd^ mutations on the binding of Vg and Yki. The Sd mutant proteins were immobilized on sensor chips and the affinity of Vg^263-382^ and Yki^30-146^ was determined by Surface Plasmon Resonance. The average and standard error of n ≥ 2 separate experiments are given. K_d_^KT^: K_d_ measured by kinetic titration. ΔΔG = ΔG_mut_ – ΔG_wt_ with ΔG_wt_ calculated from the affinity of Vg^263-382^ and Yki^30-146^ for wt^Sd^ (Table 1). ΔG = 1.986*298*LnK_d_. SE_ΔΔG_ = (SE_ΔGmut_^2^ + SE_ΔGwt_^2^)^1/2^.

|  | Vg<sup>263-382</sup> | Yki<sup>30-146</sup> |
| --- | --- | --- |
| **Asp276Ala<sup>Sd</sup>** | K<sub>d</sub><sup>KT</sup> = 5.5 ± 0.2 nM<br>ΔΔG = 1.58 ± 0.03 kcal/mol | K<sub>d</sub> = 5000 ± 300 nM<br>ΔΔG = 3.6 ± 0.1 kcal/mol |
| **Tyr435His<sup>Sd</sup>** | K<sub>d</sub><sup>KT</sup> = 24 ± 1 nM<br>ΔΔG = 2.44 ± 0.03 kcal/mol | K<sub>d</sub> = 13000 ± 300 nM<br>ΔΔG = 4.11 ± 0.04 kcal/mol |

### Presence of an Ω-loop in other proteins of the Vg/VGLL1-3 subfamily

The presence of an Ω-loop in the Sd-binding domain of Vg from *D. melanogaster* could be specific for this species or genus. In agreement with the initial observation made by Ohde et al. (Ohde et al., 2009), our analysis shows that the Vg proteins of other species belonging to the same or different insect orders contain a sequence that has strong homology with the residues located in the Ω-loop of Vg from *D. melanogaster* (Fig S7). Therefore, Vg may not be the only member of the Vg/VGLL1-3 family to possess an Ω-loop. This prompted us to turn our attention to the VGLL proteins of higher animal species and more particularly to human VGLL1-3. As previously reported (Simon et al., 2016), the β-strand:α-helix region is very well conserved between Vg and VGLL1-3 (Fig 1B). VGLL1 has no homology with Vg^323-334^ (Fig 1B) indicating that it does not possess an Ω-loop in agreement with the published structural data (Pobbati et al., 2012). VGLL3^140-151^ shows some level of homology with Vg^323-334^, but several residues (e.g. Met324^Vg^ or Arg327^Vg^) required for the interaction between Vg and Sd are missing in VGLL3 (Fig 1B) suggesting that VGLL3^140-151^ should only bind very weakly (if at all) to TEAD. In contrast, VGLL2^135-146^ shows significant sequence homology with Vg^323-334^ (Fig 1B) indicating that this region of VGLL2 may form an Ω-loop upon binding to TEAD. To measure the affinity for TEAD4 of VGLL2^135-149^, VGLL3^140-154^, YAP^85-99^ (Ω-loop) and Vg^323-337^ (Fig 1B), we used a TR-FRET assay (Hau et al., 2013, Mesrouze et al., 2014) because VGLL2^135-149^ and VGLL3^140-154^ at high concentration bind in a non-specific manner to SPR sensor chips. In the TR-FRET assay, we found that VGLL3^140-154^ has a very weak potency (IC_50_ > 500 μM; Table 3) while the IC_50_ of VGLL2^135-149^ (133 μM; Table 3) is similar to that of YAP^85-99^ (98 μM; Table 3) and Vg^323-337^ (157 μM; Table 3). This finding suggests that VGLL2, in contrast to VGLL1 and VGLL3, possesses an Ω-loop. Since VGLL2-derived peptides are rather unstable in solution (presence of three cysteines in their sequence), making them difficult to handle in crystallization experiments, we built up a molecular model of the VGLL2:TEAD complex. This model suggests VGLL2 to form an Ω-loop upon binding to TEAD, and the superimposition on the Vg:Sd structure shows that it could bind to TEAD4 in a manner similar to the way Vg binds to Sd (Fig 3). The main structural difference between bound Vg and VGLL2 resides in the linker region connecting the β-strand:α-helix and the Ω-loop. As the linker is much longer in VGLL2 (27 residues instead of 12), it adopts a more extended conformation in the VGLL2:TEAD4 complex. On the basis of these observations, we next measured the affinity of VGLL2^85-108^ (β-strand:α-helix) and VGLL2^85-149^ (β-strand:α-helix:Ω-loop) (Fig 1B) to TEAD4. VGLL2^85-108^ has an affinity (590 nM, Table 3) similar to that of its Vg counterpart, Vg^288-311^ (291 nM, Table 1). The K_d_ of VGLL2^85-108^ is in the low nanomolar range (34 nM, Table 3), showing that the presence of the Ω-loop substantially increases complex formation between VGLL2 and TEAD4. However, VGLL2^85-149^ binds less tightly to TEAD4 (Table 3) than Vg^288-327^ to Sd or TEAD4 (Tables 1 and 3). Since the isolated β-strand:α-helix and Ω-loop regions of VGLL2 and Vg have a similar affinity for TEAD or Sd, we hypothesized that the longer linker region connecting these elements in VGLL2 (Fig 1B and 3) has a negative impact on the interaction with TEAD4. To test this hypothesis, we designed a hybrid peptide (hereafter referred to as Flyman) formed of the β-strand:α-helix and Ω-loop of VGLL2 connected by the linker from Vg (Fig 1B). Flyman is more potent than VGLL2^85-149^ and its potency is similar to that of Vg^288-337^ (Table 3) suggesting that the linker found in VGLL2 has a negative effect on the binding of VGLL2^85-149^ to TEAD4.

**Figure 3.**
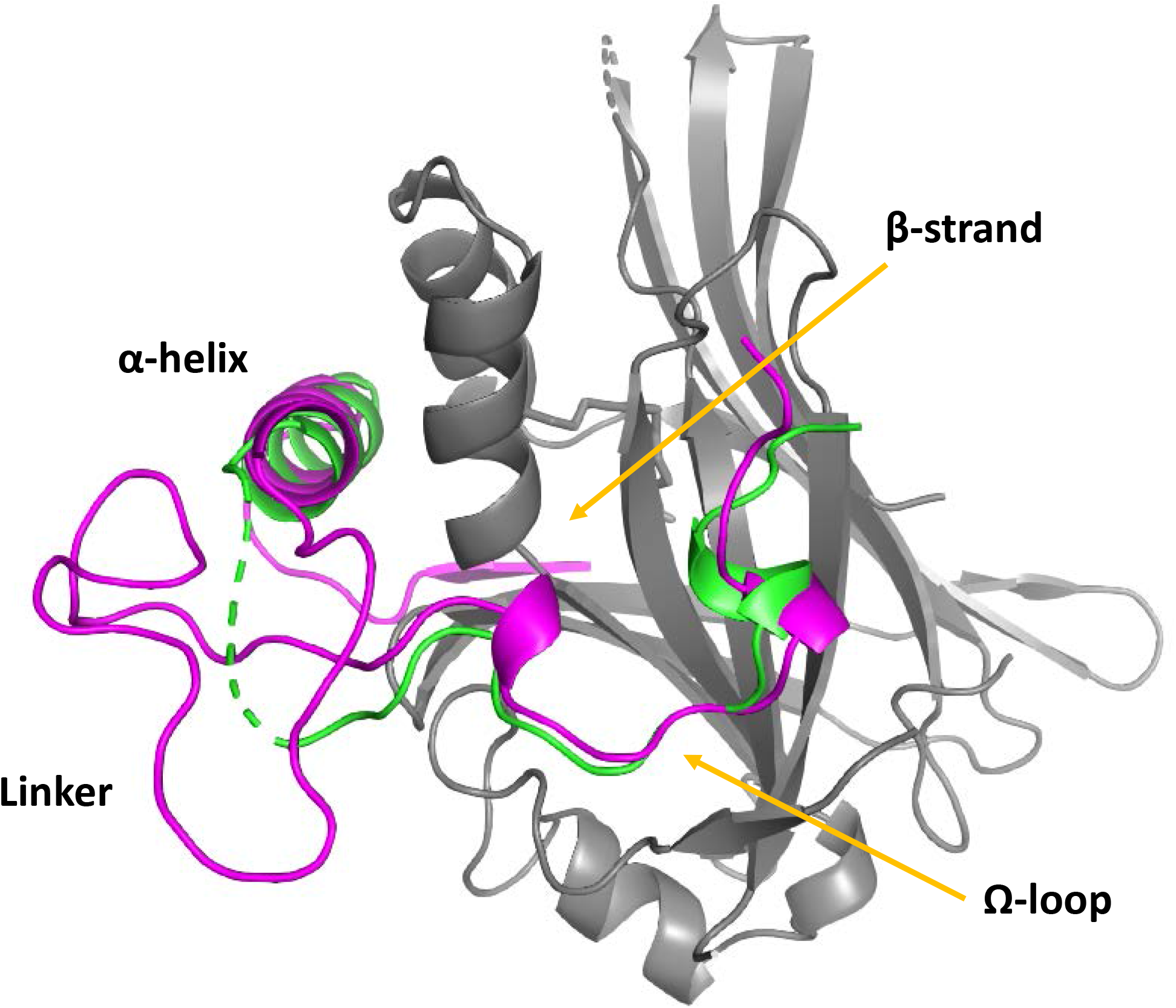
Molecular model of the VGLL2:TEAD4 complex. A molecular model of the VGLL2^85-149^:TEAD4^217-434^ complex was superimposed on that of the crystal structure of the Vg^298-337^:wt^Sd^ complex (PDB 6Y20). VGLL2^85-149^, Vg^298-337^ and TEAD4^217-434^ are represented in magenta, green and grey, respectively. The secondary structure elements of VGLL2^85-149^ are indicated. The figure was drawn with PyMOL (Schrödinger Inc., Cambridge, MA).

**Table 3.** Potency of VGLL2 and VGLL3 derived peptides. The potency of the peptides (IC_50_) was measured in a TR-FRET assay. The K_d_ values were determined at equilibrium by Surface Plasmon Resonance. The average and standard error of n ≥ 2 separate experiments are given. ^a^. IC_50_s below the assay limit fixed by the concentration of TEAD4 present in the FRET assay. n.d. not determined.

|  | YAP<sup>85-99</sup> | Vg<sup>323-337</sup> | VGLL2<sup>135-149</sup> | VGLL3<sup>140-154</sup> | VGLL2<sup>85-108</sup> | VGLL2<sup>85-149</sup> | Vg<sup>288-337</sup> | Flyman |
| --- | --- | --- | --- | --- | --- | --- | --- | --- |
| **IC<sub>50</sub> (nM)** | 98000<br>± 2000 | 160000<br>± 11000 | 133000<br>± 7000 | > 500000 | 340<br>± 30 | 6.3<br>± 0.4 | a | a |
| **K<sub>d</sub> (nM)** | 58000<br>± 800 | n.d. | n.d. | n.d. | 590<br>± 10 | 34<br>± 4 | 3.1<br>± 0.2 | 4.3<br>± 0.2 |

To determine whether the Ω-loop region of VGLL2, VGLL2^135-146^ also contributes to the interaction with TEAD in cells in the context of the full-length proteins, N-terminally V5-tagged wt^TEAD4^ and (FLAG)3-wt^VGLL2^ were co-transfected into HEK293FT cells harboring a genomic deletion of YAP and TAZ (transcriptional co-activator with PDZ-binding motif, a paralog of YAP) (see below). Following V5-mediated immunoprecipitation, wt^TEAD4^ and bound wt^VGLL2^ were monitored using Western blot with V5 and FLAG-antibodies, respectively. wt^VGLL2^ was efficiently co-immunoprecipitated with wt^TEAD4^, while it was not detected in similar experiments carried out in the absence of V5-tagged TEAD4 (empty vector control) (Fig 4A). To test whether VGLL2 interacts with TEAD4 via its Ω-loop region, we used the delΩ^VGLL2^ mutant protein, in which the residues 135-146 have been deleted. The deletion significantly decreases the amount of VGLL2 protein co-immunoprecipitated with wt^TEAD4^ indicating that the region 135-146 is required for efficient binding to TEAD4 (Fig 4A). To determine whether VGLL2 interacts with TEAD4 via its Ω-loop binding pocket, we used the Asp272Ala^TEAD4^ mutant. According to our molecular model, this mutation should prevent the formation of a salt bridge with Arg139^VGLL2^ and it should have the same disruptive effect on the VGLL2:TEAD4 complex as the Asp246Ala^Sd^ mutation on the Vg:Sd complex (Table 2). The amount of wt^VGLL2^ co-immunoprecipitated with Asp272Ala^TEAD4^ is lower than that detected in the presence of wt^TEAD4^ (Fig 4A). Altogether, these findings show that the Ω-loop region, VGLL2^135-146^, is required for an efficient interaction between full-length VGLL2 and TEAD4 in cells. Upon quantification of the immunoprecipitated fraction (IP) over input ratio across three separate experiments which exhibited a similar pattern, the Asp272Ala^TEAD4^ mutation or the Ω-loop deletion of VGLL2 led to a 2 to 2.5-fold reduction in the VGLL2:TEAD4 interaction (Fig 4B). The combination of these two alterations (Fig 4A) did not further decrease the extent of co-immunoprecipitation (~ 2-fold reduction, Fig 4B). This supports the notion that the two mutations affect the same binding interface; if they disrupted interactions at distinct interfaces, then an additive or a synergistic effect would be expected.

**Figure 4.**
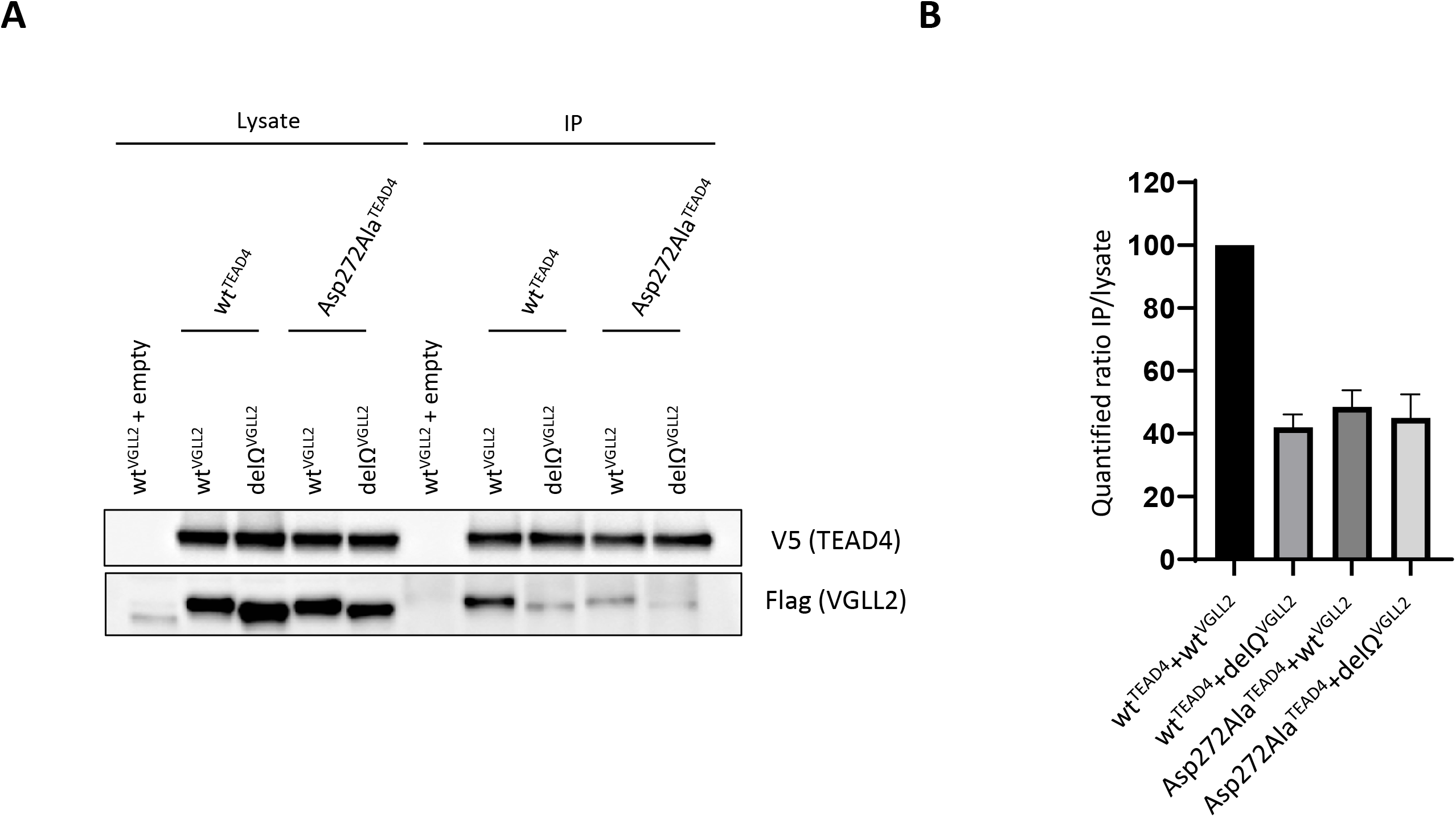
Interaction between VGLL2 and TEAD4 in cells. **A.** V5-tagged wt^TEAD4^ or Asp272Ala^TEAD4^ constructs were co-transfected with (FLAG)3-tagged wt^VGLL2^ or delΩ^VGLL2^ in the indicated combinations into HEK293FT double YAP/TAZ knockout cells. TEAD4 was immunoprecipitated with a V5-specific antibody and co-immunoprecipitated VGLL2 was detected by anti-FLAG Western blot (left panel). Empty: Empty vector control. IP: Immunoprecipitated fraction. **B.** Image J-based (https://imagej.nih.gov/ij/) quantification (average and SD of three separate experiments) was completed on the Western blot signals of indicated co-IP combinations, and IP/lysate ratios are shown relative to the wt^TEAD4^/wt^Vgll2^ interaction set to 100% (right panel). The data presented correspond to the average and standard deviation of three separate experiments.

Similar results with regard to VGLL2:TEAD4 complex formation and the impact of the Asp272Ala^TEAD4^ and delΩ^VGLL2^ mutations were observed in wild-type HEK293FT (i.e. with intact YAP and TAZ). However, the inter-experimental reproducibility across the four probed co-IP conditions was less robust. We hypothesized that this was a consequence of the competition between YAP, TAZ and VGLL2, which bind with different affinities to the Ω-loop binding pocket of TEAD, leading to an increased susceptibility to multi-parametric experimental variations in complex stoichiometry. To facilitate the mechanistic dissection and increase experimental robustness, we therefore opted for the use of double YAP/TAZ knockout cells in our studies. We also repeatedly observed that the expression levels of FLAG-tagged VGLL2 were significantly higher in co-transfection experiments performed with TEAD4 proteins (Fig 4). In contrast to our previously reported observations for FAM181A (Bokhovchuk, Mesrouze et al., 2020), this effect happened with wild-type and mutant forms of TEAD4 and VGLL2, suggesting a the formation of a complex formation between these two proteins. This is presumably mediated through the interactions taking place at the β-strand:α-helix binding site, which are unaffected by the Asp272Ala^TEAD4^ or the deletion of the Ω-loop and may enhance the stability of co-overexpressed VGLL2 in this experimental setting.

### Functional role of the Ω-loop from Vg in Drosophila

To investigate the functional contribution of the Ω-loop in an *in vivo* context, we generated several transgenic drosophila fly lines that conditionally express different HA-tagged forms of Vg. We obtained UAS lines containing cDNAs encoding either full-length Vg, or mutant versions bearing a deletion of the Ω-loop (delΩ), a deletion of the β-strand:α-helix (delβ:α) or the deletion of the whole Sd-binding domain of Vg (delβ:α:Ω). In the larval wing primordium, the wing imaginal disc, Vg-Sd interaction plays a major role in the specification of the wing cell fate; the deletion of Vg leads to a complete loss of the wing pouch and subsequently the adult wing (Williams, Bell et al., 1991). Strikingly, ectopic expression of Vg outside its wild-type expression pattern leads to ectopic wing outgrowths in the wing imaginal disc, whereas ectopic expression in other imaginal discs leads to their transformation into wing fate (Baena-Lopez & Garcia-Bellido, 2003, Simmonds et al., 1998). We used such *in vivo* ectopic expression assays to probe the function of the different forms of Vg mentioned above.

Upon 24 h ectopic expression using an en-gal4 driver line expressed in the posterior compartment of the wing imaginal disc, we detected comparable level of expression of the four HA::Vg proteins (Fig 5A). Expression of HA::Vg or HA::Vg^delΩ^ resulted in tissue outgrowth and repression of the hinge specific protein Homothorax (Hth) in the posterior compartment (Fig 5B), suggesting hinge to wing transformation via these two functional Vg proteins. Expression of HA::Vg^delα:β^ and HA:Vg^delα:β:Ω^ did not lead to morphological defects and no change in the distribution of Hth were observed. Consistently, the presumptive wing region, delimited by a Wingless (Wg) ring pattern, was expanded upon expression of HA::Vg or HA::Vg^delΩ^, but not when HA::Vg^delα:β^ and HA:Vg^delα:β:Ω^ where expressed (Fig 5C). These results show a strong dependency on the β-strand and α-helix for full Vg function, but fail to reveal differences between wild-type Vg and Vg^delΩ^.

**Figure 5.**
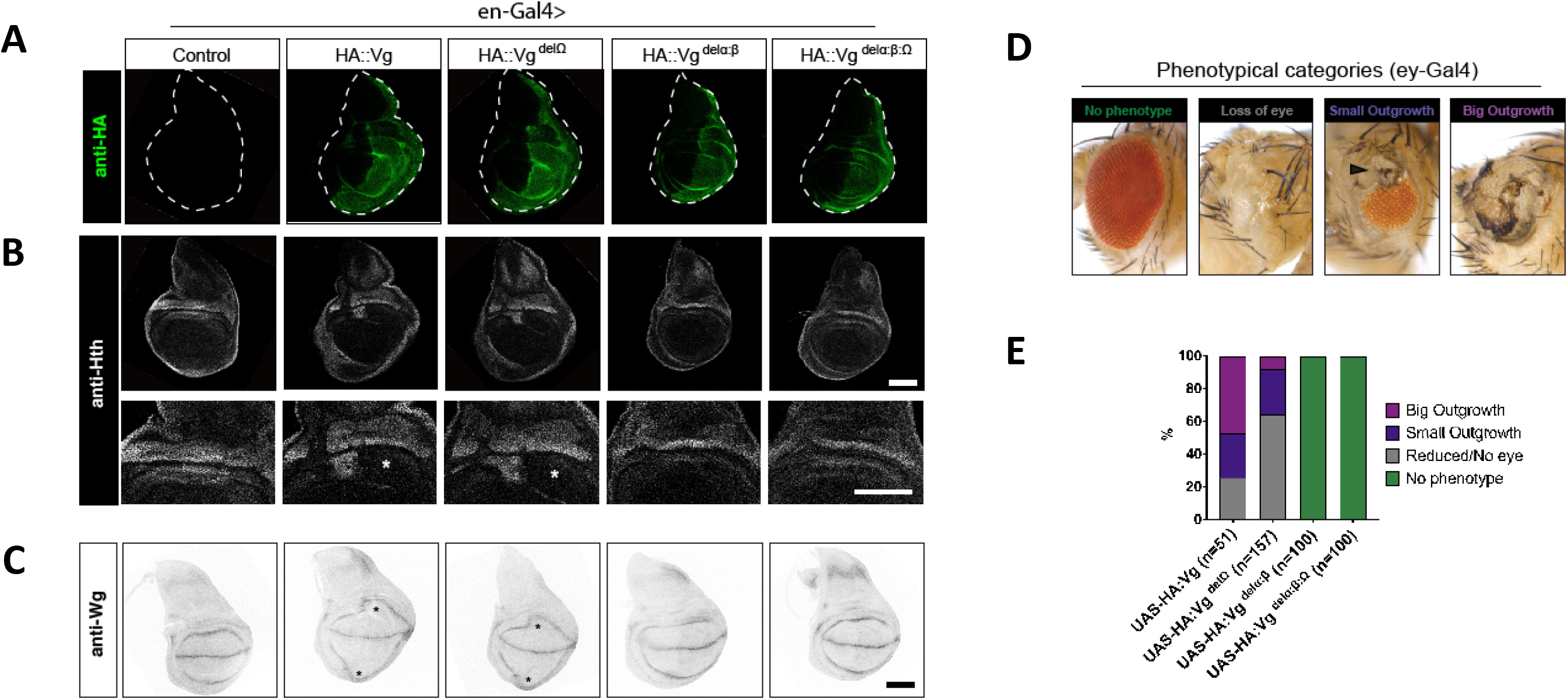
Functional characterization of Vg mutant proteins. **A.** Immunostaining of N-terminal tagged HA::Vg constructs upon 24 h expression with an en-Gal4 driver, active in posterior compartment cells. Dashed line delimits the wing disc. **B.** Hth immunostaining of same discs as in **A**. HA::Vg and HA::Vg^delΩ^ expressing discs show a disruption of the Hth localization pattern accompanied with tissue outgrowth. Asterisks point the affected area. Scalebars 100 μm. **C.** Wg localization pattern in disc of same genotype as in **A** and **B**. Asterisks show expansion of presumptive wing tissue in HA::Vg and HA::Vg^delΩ^ discs. **D.** Phenotypical classes scored upon expression of the Vg constructs with the ey-Gal4 driver. Arrowhead points the small outgrowth and arrow points to the large outgrowth. E. Quantification of the observed phenotypes.

Therefore, we turned our attention to a different ectopic expression assay and expressed the four Vg constructs under the eye-specific driver line ey-Gal4. As expected based on previous studies (Takanaka & Courey, 2005), expression of HA::Vg resulted in different eye phenotypes (Fig 5B), including wing-like tissue outgrowths in the eye, with a high phenotypic penetrance (Fig 5C). Expression of HA::Vg^delΩ^ was also able to generate such outgrowths, but the size and penetrance of these was lower than upon HA::Vg expression. Expression of HA::Vg^delα:β^ and HA::Vg^delα:β:ω^ failed to change the fate of eye cells. Together, these results show again a strong dependence on the TONDU domain (β-strand:α-helix region), and also reveal a moderate but significant role of the Ω-loop in the function of Vg.

## Discussion

VGLL1 and YAP bind via a similar β-strand:α-helix motif to an overlapping region at the surface of the TEAD transcription factors (Chen et al., 2010, Li et al., 2010, Pobbati et al., 2012). This motif, which is highly conserved amongst the Vg/VGLL1-3 proteins (TONDU domain) (Simon et al., 2016), is the only structural element known to be involved in the interaction between these co-factors and the TEAD transcription factors. However, YAP (and its paralog TAZ (Kaan, Sim et al., 2017)) requires in addition to the β-strand:α-helix motif an Ω-loop for efficient binding to TEAD (Hau et al., 2013, Li et al., 2010). Therefore, the main difference between the TEAD-binding domain of the Vg/VGLL1-3 and YAP-like proteins is the presence of an Ω-loop in the latter (Gibault, Coevoet et al., 2018, Pobbati et al., 2012, Santucci et al., 2015). The data presented in this article challenge this view, revealing that the Sd/TEAD-binding domain of some Vg/VGLL1-3 proteins also contains an Ω-loop. We demonstrate that Vg from *D. melanogaster* has an Ω-loop that is required for tight binding to Sd and our *in vivo* assays show that in some ectopic assays (such as the transformation of the eye to wing-like structures), full function of the Vg protein requires the Ω-loop. Furthermore, we show that human VGLL2, in contrast to human VGLL1 and VGLL3, also possesses an Ω-loop that contributes to its interaction with TEAD. Altogether, these findings indicate that the Sd/TEAD-binding domain of the Vg/VGLL1-3 proteins exists in two different forms: a β-strand:α-helix motif or a β-strand:α-helix:Ω-loop motif. This suggests that to be fully functional, all the Vg/VGLL1-3 proteins must possess a TONDU domain but that the presence of an Ω-loop is not required for some of them (VGLL1, 3) while for others (Vg, VGLL2) it might be needed to reach complete activity. In the following, we shall discuss a hypothesis to explain the evolution of this protein family.

The whole genome duplications that occurred at the origin of the vertebrate lineage (Dehal & Boore, 2005, Panopoulou, Hennig et al., 2003) probably led to the presence of duplicates of an ancestor *Vg* gene in the genome of early vertebrates. As VGLL1-3 from vertebrates derive from *Drosophila* Vg (Faucheux, Naye et al., 2010), it is likely that the duplicated VGLL proteins present in early vertebrates contained an Ω-loop. The presence of multiple forms of the same gene in genomes enables the accumulation of mutations that can lead to the modification of the ancestral gene (Huminiecki & Wolfe, 2004, Lynch & Katju, 2004, Seoighe, Johnston et al., 2003). The nanomolar affinity of the isolated TONDU domain of Vg/VGLL2 and the triple-digit micromolar affinity of their isolated Ω-loop indicate that the TONDU domain is the “hot spot” for the interaction of these proteins with Sd/TEAD. Since isolated TONDU domains and the TEAD-binding domain of Yki/YAP (or TAZ) have a similar affinity for TEAD (here and (Bokhovchuk et al., 2019)), the VGLL proteins need to contain only a TONDU domain to bind to TEAD and to compete with YAP (or TAZ) for binding to TEAD, as is observed in cells with VGLL1 and VGLL3 (Figeac, Mohamed et al., 2019, Pobbati et al., 2012). Therefore, mutations in the Ω-loop region of the VGLL proteins from early vertebrates could take place and be tolerated during evolution because the mutated proteins would still be able to bind with sufficient affinity to TEAD via their TONDU domain. We propose that the accumulation of mutations during this evolutionary process led to the loss of the Ω-loop in VGLL1 and VGLL3. VGLL2 followed a different trajectory since it still contains a functional Ω-loop. Nonetheless, the linker connecting this Ω-loop to the TONDU domain has a negative contribution to the interaction with TEAD. It is difficult to determine whether the ancestral VGLL variant leading to VGLL2 had a similar linker or whether mutations have progressively transformed it from a shorter Vg-like linker to a longer one. In the second case, the transformation of the linker could be the first step in leading to the loss of the Ω-loop and VGLL2 could be an evolutionary intermediate between an ancestral VGLL protein and VGLL1/VGLL3.

The duplication of genes and their subsequent transformation by mutations is a well-accepted concept in the literature, and the Vg/VGLL1-3 and Yki/YAP families provide a beautiful example of how structural and functional requirements have contributed to the evolution of two co-factor families that regulate the same transcription factors through interaction with an overlapping binding site. The loss of the β-strand:α-helix motif dramatically decreases the affinity of Vg/VGLL1-3 (“hot spot” loss) while in Yki/YAP it prevents the ability to compete with Vg/VGLL1-3 and reduces affinity. Therefore, this motif must be preserved in both families. The loss of the Ω-loop generates Vg/VGLL1-3 variants that are still able to bind to Sd/TEAD and to compete with Yki/YAP, while in the latter it abolishes binding (“hot spot” loss). Therefore, the Vg/VGLL1-3 proteins but not the Yki/YAP proteins can lose their Ω-loop. These structural and functional constraints may explain why a β-strand:α-helix or a β-strand:α-helix:Ω-loop motif is present in the Sd/TEAD-binding domain of the Vg/VGLL1-3 proteins while, to our knowledge, only a β-strand:α-helix:Ω-loop motif is found in the Yki/YAP proteins.

## Material and methods

### Peptides

The synthetic peptides (both N-acetylated and C-amidated) were purchased from Biosynthan (Germany). The purity (>90%) and the chemical integrity of the peptides were determined by LC-MS from 10 mM stock solutions in 90:10 (v/v) DMSO:water. VGLL2^85-149^, which contains three cysteines (Fig 1B), was dissolved in 50:50 (v/v) acetontrile:(water+1mM TCEP) to avoid oxidation mechanisms induced by DMSO. VGLL2^85-108^ was also prepared in the same solvent. The experiments conducted with VGLL2^85-149^ were done with freshly made solutions (less than 72 h old). The peptide solutions were stored at −20°C and for each experiment the solutions were centrifuged to remove potential aggregates; the supernatant was dosed by HPLC and the integrity of the peptide was evaluated by LC-MS. To determine whether acetonitrile affects the output of the biochemical assays, Vg^288-337^ was also dissolved in 50:50 (v/v) acetonitrile:(water+1mM TCEP). The K_d_ value of this peptide preparation – 4.1 ± 0.1 nM (SPR) – is similar to that obtained with a DMSO solution of peptide (3.1 nM; Table 3) showing that the acetonitrile (2% final concentration) does not interfere with our assays.

### Cloning, expression and purification of the proteins for the biochemical and biophysical assays

The amino acid sequences of Scalloped (Sd, UniProt P30052, amino acids 223-440), Yorkie (Yki, UniProt Q45VV3, amino acids 30-146), and Vestigial (Vg, UniProt Q26366, amino acids 263-382) from *Drosophila melanogaster* were back-translated into an *Escherichia coli* codon-optimized DNA sequence by the GeneArt online ordering tool and synthesized by GeneArt (Thermo Fisher, Switzerland). The DNA fragment encoding Sd^223-440^ was PCR amplified and inserted into a pET28-derived vector providing an N-terminal His-HRV3C-Avi-tag by HiFi DNA Assembly Cloning (New England Biolabs, Ipswich, MA) according to the instructions of the manufacturer. The mutations to generate Asp276Ala^Sd^ and Tyr435His^Sd^ were introduced by the QuikChange II Lightning Site-Directed Mutagenesis kit (Agilent, Santa Clara, CA) according to the manufacturer’s instructions. The DNA fragments encoding Yki^30-146^ and Vg^263-382^ were PCR amplified and cloned as described above into a pET derived vector providing an N-terminal His-tag followed by an HRV3C protease site. In a second step, the coding sequences for strep-tagII and rubredoxin were inserted into both constructs between the His-tag and the HRV3C site by HiFi DNA Assembly Cloning. All expression constructs were confirmed by Sanger sequencing. Yki^30-146^ and Vg^263-382^ were expressed in *E. coli* BL21(DE3) cells (Novagen, Madison, WI). For the expression of different Sd proteins, the *E. coli* BL21(DE3) cells contained in addition a plasmid encoding for biotin ligase BirA (acetyl-CoA carboxylase biotin holoenzyme synthetase). Transformed cells were diluted in LB medium containing 25 μg/ml kanamycin (Invitrogen, Carlsbad, CA) and in addition 34 μg/ml chloramphenicol (Sigma-Aldrich, St Louis, MI) for Sd expression. Pre-cultures were incubated overnight at 37°C with constant agitation and diluted into fresh medium containing antibiotics and the cultures were grown at 37°C until OD600 = 0.8. Protein expression was induced by adding 1 mM isopropyl β-D-1-thiogalactopyranoside (AppliChem GmbH, Germany) (and 135 μg/ml biotin (Sigma-Aldrich, St Louis, MI) for Sd expression). The cells were grown overnight at 18°C, harvested by centrifugation, frozen in dry ice and stored at −80°C. All purification steps were carried out at 4-8°C unless stated otherwise. Frozen cell pellets were thawed on ice and resuspended in Buffer A (50 mM TRIS·HCl pH 8.0, 1 M NaCl, 2 mM MgCl_2_, 1 mM Tris(2-carboxyethyl)phosphine (TCEP), 20 mM imidazole, 0.1% (v:v) Tween 20, 0.5 mM biotin (Sigma-Aldrich, St Louis, MI), 5% (v:v) glycerol), cOmplete EDTA free protease inhibitor (1 tablet/50 ml buffer; Roche, Switzerland), and TurboNuclease (20 μl/50 ml buffer; Sigma-Aldrich, St Louis, MI). The cells were disrupted by 3 passes through an EmulsiFlex C3 high pressure homogenizer (Avestin Inc., Canada) at 800–1000 bar and centrifuged at 43000 x g for 30 min. The cleared supernatant was loaded onto two HisTrap HP 1 mL columns (GE Healthcare, Chicago, IL) mounted in series and equilibrated with Buffer A. The columns were washed with Buffer A followed by Buffer B (50 mM TRIS·HCl pH 8.5, 100 mM NaCl, 2 mM MgCl_2_, 1 mM TCEP, 20 mM imidazole, 5% (v:v) glycerol). Bound proteins were eluted with a linear gradient of Buffer B containing 250 mM imidazole. The fractions containing Scalloped were pooled and the His6-tag was cleaved off by overnight incubation during dialysis in Buffer B with 1% (w:w) recombinant HRV 3C protease (produced in-house). The dialysed protein was loaded onto two His-Trap HP 1 mL columns (GE Healthcare, Chicago, IL) mounted in series and equilibrated with Buffer B. The flow-through fractions containing the cleaved protein were collected, pooled, and concentrated by centrifugation in Amicon Ultra-15 centrifugal units (MilliporeSigma, Burlington, MA). After determination of the protein concentration by RP-HPLC, a threefold molar equivalent of myristoyl coenzyme A (lithium salt; Sigma-Aldrich, St Louis, MI) was added and incubated at room temperature for 30-60 min. The reaction products were loaded onto a HiLoad 16/600 Superdex 75 prep grade column (GE Healthcare, Chicago, IL) equilibrated with 50 mM TRIS·HCl pH 8.0, 100 mM NaCl, 2 mM MgCl_2_, 1 mM TCEP, 5% (v:v) glycerol. Protein fractions containing Scalloped were concentrated by centrifugation in Amicon Ultra-15 cartridges (MilliporeSigma, Burlington, MA), aliquoted, snap-frozen in dry ice and stored at −80°C. Yki^30-146^ and Vg^263-382^ were expressed as described for Scalloped except that *E. coli* NiCo21 (DE3) cells (New England Biolabs, Ipswich, MA) grown in TBmod plus MOPS medium supplemented with 25μg/ml kanamycin and no biotin were used. Yki^30-146^ was purified as described for Scalloped but 300 mM NaCl was used during protease cleavage. For purification of Vg^263-382^, Buffer B contained during protease cleavage 300 mM NaCl and 0.1% (v/v) CHAPS (Sigma-Aldrich, St Louis, MI). The HiLoad 16/600 Superdex 75 prep grade column (GE Healthcare, Chicago, IL) was either equilibrated as for the other proteins or alternatively equilibrated with 50 mM HEPES, 100 mM KCl, 0.25 mM TCEP, 1 mM EDTA, 0.05% Tween 20, pH 7.4. The purity, quantity and identity of all proteins were determined by RP-HPLC and LC-MS (Fig S3).

### Circular Dichroism, Surface Plasmon Resonance and FRET assay

Representative experiments and methodology are presented on: Fig S4, Circular Dichroism; Fig S5, Surface Plasmon Resonance; Fig S8, TR-FRET assay.

### Structural biology

The DNA fragment encoding for wt^Sd^, obtained as described above, was PCR amplified and inserted into a pET28-derived vector providing an N-terminal His-HRV3C-tag by HiFi DNA Assembly Cloning (New England Biolabs, Ipswich, MA) according to the instructions of the manufacturer. The construct encoding for wt^Sd^ was expressed in *E. coli* BL21(DE3) cells (Novagen, Madison, WI) without addition of biotin to the medium and purified according the protocol identical to the one described above for the biotinylated Sd proteins. Crystals of the complex between untagged wt^Sd^ and Vg^298-337^ were grown at 293°K using the sitting drop vapour diffusion method. wt^Sd^ (8.3 mg/ml) in 50 mM Tris pH 8.0, 250 mM NaCl, 2 mM MgCl_2_, 1 mM TCEP and 5% glycerol was pre-incubated with 0.5 mM Vg^298-337^ (molar ratio Vg^298-337^ / wt^Sd^ ~1.5). For crystallization, the peptide-protein complex was mixed with an equal volume of the reservoir solution (0.3 μL + 0.3 μL). Initial crystals were obtained using 0.1 M Bis-Tris pH 6.5 and 25% PEG3350 as reservoir solution. Micro seeds were prepared from these conditions and used for further crystallization trials. Diffraction quality crystals were obtained using 0.2 M MgCl_2_,6H_2_O, 0.1 M Bis-Tris pH 6.5 and 25% PEG3350 as reservoir solution. Prior to shock cooling in liquid nitrogen, the crystals were soaked for a few seconds in reservoir solution containing 30% glycerol. X-ray diffraction data were collected at the Swiss Light Source (SLS, beamline X10SA) using an Eiger pixel detector. Raw diffraction data from two crystals originating from the same crystallization drop were analysed and processed using the autoPROC (Vonrhein, Flensburg et al., 2011) / STARANISO (Tickle et al., Global Phasing Ltd., Cambridge, UK) toolbox. The structure was solved by molecular replacement with PHASER (McCoy, Grosse-Kunstleve et al., 2007) using the coordinates of previously solved in-house structures of TEAD4 as the search model. The software programs COOT (Emsley & Cowtan, 2004) and BUSTER (Bricogne et al. Buster-TNT 2.X, Global Phasing Ltd., Cambridge, UK) were used for iterative rounds of model building and structure refinement.

### Molecular modelling

A model of TEAD4^217-434^ in complex with VGLL2^85-149^ was constructed by homology. The crystal structure of YAP in complex with TEAD4 (PDB 6GE3) was used as the starting point. The regions corresponding to the α-helix and the linker region of YAP were deleted and the Ω-loop of YAP, YAP^85-99^, was mutated to mimic the Ω-loop of VGLL2, VGLL2^135-149^. The crystal structure of TEAD4 in complex with VGLL1 (PDB 5Z2Q) was next superimposed on this partial model. TEAD4 from this structure was deleted and the β-strand:α-helix region of VGLL1, VGLL1^27-51^, was mutated to mimic VGLL2^85-108^. Finally, an arbitrary conformation allowing the α-helix and the Ω-loop sequences to be connected in 3D was given to an amino acid stretch corresponding to the linker region VGLL2^109-136^. Molecular dynamics simulation of 10 ns with an explicit water solvation model was run using the Desmond module (default parameters) in the molecular modelling package Maestro (Schrödinger Inc. Cambridge, MA). The β-strand:α-helix and Ω-loop regions of VGLL2 were quite stable during the simulation, maintaining their secondary structure. At the end of the simulation, the linker region converged towards a loop conformation with no regular secondary structure that appeared stabilized by an aromatic stacking interaction between residues Tyr115^VGLL2^ and Phe196^VGLL2^. *Cellular biology.* HEK293FT YAP/TAZ double knockout (ko) cells were established as follows: The sgRNA (CRISPR) targeting sequences for YAP (‘#1’: *TGG GGG CTG TGA CGT TCA TC*) and WWTR1 (‘#2’: *TCC AGC ACC GAC TCG GG* and ‘#B’: *GTC CAG CAC CGA CTC GTC GG*) were cloned into pU6gRNA_CMV-SPyCas9-T2Apuro. 293FT YAP ko cells were created by transfecting pU6gRNA_YAP_1_CMV-SPyCas9-T2Apuro in 293FT cells using Lipofectamine 2000 (Thermo Fisher, Switzerland). Following transient puromycin selection, clones were picked and characterized for (homozygous) absence of YAP at genomic and protein expression level. Selected 293FT YAP ko clones were subsequently co-transfected with pU6gRNA_WWTR1_2_CMV-SPyCas9-T2Apuro and pU6gRNA_WWTR1_B_CMV-SPyCas9-T2Apuro. Following the same selection procedure, the picked clones were characterized for (homozygous) absence of TAZ (WWTR1) at genomic and protein expression level, resulting in verified HEK293FT double YAP/TAZ ko cells. VGLL2 cloning was performed as follows: VGLL2 (RefSeq NM_182645.3) was amplified by standard PCR from a pDONR221-VGLL2 (human codon-optimized) clone, obtained from GeneArt (Germany), using primers containing N-terminal (FLAG)3 epitope. The PCR product was cloned by Gateway reaction into pcDNA3.1 Hygro-DEST (Invitrogen, Carlsbad, CA), according to the manufacturer’s protocol. The pcDNA3.1 Hygro-DEST_VGLL2_Ω_loop mutant was generated by use of the QuikChange Lightning Site-Directed Mutagenesis kit (Stratagene, San Diego, CA), with oligo 5’*CCC TTG GAG AGA CTG CAG CTT CTA TCA GGC TCC TGT GCC* 3’ (removing amino acid positions 134-149) and the sequence of resulting cDNA clones verified. All other cellular biology tools and methods, namely TEAD4 constructs and coimmunoprecipitations, were essentially as described previously (Bokhovchuk et al., 2020, Mesrouze et al., 2017a).

### In vivo biology

Molecular cloning and generation of transgenic fly lines. pUAS-HA:Vg-attb constructs were generated by Gibson Assembly from amplified fragments of the Vg cDNA and the pUASt-attb vector. A GGGS linker was added between HA and Vg ORF to reduced possible positional effect. The resulting constructs (pUAS-HA::Vg, pUAS-HA::Vg^delΩ^ (lacking amino acids 323 to 337), pUAS-HA::Vg^delα:β^ (lacking amino acids 288 to 311) and pUAS-HA::Vg^delα:β:Ω^ (lacking amino acids 288 to 337)) were injected into eggs laid by vasa-integrase; attB86Fb flies to establish the transgenic lines. Fly stocks and fly husbandry. en-Gal4 and ey-Gal4 (2^nd^ Chromosome) are described in Flybase. For temporal control of Gal4 function, animals carrying the en-Gal4, UAS-HA::Vg and the tubulin-Gal80ts insertions were kept at 17°C during development and transferred at 29°C for 24 h before dissection. Immunostaining and imaging. The primary antibodies used in this study were: 1:300 anti-HA (3F10, Roche, Switzerland) and 1:500 anti-Hth (Gift from N. Azpiazu, Autonomous University of Madrid, Spain). Goat anti-Rabbit Alexa680 and goat anti-Rat Alexa555 secondary antibodies (Thermo Fisher Scientific, Waltham, MA) were employed. The immunostaining protocol was performed as follows: third instar stage larvae were dissected in cold PBS (Cat. number 20012019, Gibco Thermo Fisher Scientific, Waltham, MA) and fixed 30 min in 4% PFA-PBS solution. After washing in PBS and permeabilizing in PBT (0.3% Triton-PBS), samples were blocked in 5% NGS-PBT for 1 h. Samples were then incubated in primary antibody diluted in blocking solution over-night. Next morning, samples were washed in PBT and incubated in secondary antibody diluted in blocking solution for 2 h. After extensive washing first with PBT and then with PBS, imaginal discs were separated from the cuticle and mounted in Vectashield with DAPI (H-1200, Vector Laboratories, Burlingame, CA). All steps were performed at roomtemperature except primary antibody incubation that was performed at 4°C. Wing disc imaging was performed with LSM point confocal 880.

## Data availability

The refined coordinates of the Vg:Sd complex structure have been deposited in the Protein Data Bank (www.wwpdb.org) under accession code 6Y20.

## Acknowledgements

The work of MA and GA was supported by the Kantons Basel-Stadt and Basel-Land and by grants from the SNSF to MA. GA was supported by a “Fellowship for Excellence” of the International PhD Program in Molecular Life Sciences of the Biozentrum, University of Basel.

## Author contributions

YM, GA, FB, TM, CD, FV, MM, CZ, PF and RW performed research and analysed data. TV, DE, PF, CS, TS, MA and PC supervised the experimental work. MA and PC wrote the manuscript. PC designed research.

