## Supplementary Information for "A new perspective on the evolution of the interaction between the Vg/VGLL1-3 proteins and the TEAD transcription factors"

**Table S1.** Crystallographic data collection and refinement statistics for the $Vg^{298\text{-}337}$:$Sd^{222\text{-}440}$ complex structure.

**Figure S1.** Structure of the YAP:TEAD and VGLL1:TEAD complexes.

**Figure S2.** Primary sequence alignment of TEAD4 and Sd.

**Figure S3.** Representative LC-MS analyses of the recombinant proteins.

**Figure S4.** Circular dichroism analysis of $wt^{Sd}$, $Asp276Ala^{Sd}$ and $Tyr435His^{Sd}$.

**Figure S5.** Surface Plasmon Resonance.

**Figure S6**. Superimposition of the structure of the $Vg^{298\text{-}337}$:$Sd^{223\text{-}440}$ complex on that of the YAP:TEAD complex.

**Figure S7**. Primary sequence alignments of Vg proteins from different insect species.

| | $Vg^{298\text{-}337}$:$Sd^{222\text{-}440}$ |
|---|---|
| **Data collection**[a] | |
| Space group | $P2_12_12_1$ |
| Cell (Å) | a=48.2, b=62.0, c=156.9 |
| Resolution (Å) | 1.85 (2.01-1.85)[b] |
| $R_{merge, all}$ (%) | 30.7 (212.3) |
| Mean I/sigI | 9.3 (1.6) |
| Completeness spherical (%) | 74.6 (16.9) |
| Completeness ellipsoidal (%) | 93.3 (50.9) |
| Multiplicity | 16.7 (16.2) |
| **Refinement**[c] | |
| Resolution (Å) | 78 - 1.85 |
| No. reflections | 30586 |
| $R_{work/}$ $R_{free}$ (%) | 19.8 / 22.4 |
| R.m.s deviations | |
| Bond lengths (Å) | 0.008 |
| Bond angles (deg) | 0.98 |
| Complex molecules in AU | 2 |
| Myristate | Covalent to $Lys350^{Sd}$ in protein chain A<br>Non-covalent in protein chain B |
| **PDB code** | 6Y20 |

**Table S1.** Crystallographic data collection and refinement statistics for the $Vg^{298\text{-}337}$:$Sd^{222\text{-}440}$ complex structure. [a]Values as reported in autoPROC (see material and methods); [b]Highest resolution shell is shown in parentheses; [c]Values as defined in BUSTER (see material and methods).

**Figure S1.** Structure of the YAP:TEAD and VGLL1:TEAD complexes. The structures of the YAP:TEAD (PDB 3KYS (Li et al., 2010)) and of the VGLL1:TEAD (PDB 4EAZ (Pobbati et al., 2012)) complexes were superimposed with PyMOL (Schrödinger Inc., Cambridge, MA). YAP, VGLL1 and TEAD are represented in green, magenta and grey, respectively. The secondary structure elements involved in the interaction with TEAD are labelled.

**Figure S2.** Primary sequence alignment of TEAD4 and Sd. The region corresponding to the YAP/VGLL-binding domain of TEAD4, $TEAD4^{217-434}$ (UniProt Q15561) and $Sd^{223-440}$ (UniProt P30052) is aligned with T-Coffee (Notredame, Higgins et al., 2000).

**Figure S3.** Representative LC-MS analyses of the recombinant proteins. The figure represents LC chromatograms. The amount of protein (in µg) used for the analyses are indicated and the mass of the protein identified in the peaks are given. The numbers in brackets correspond to the theoretical molecular weight of the corresponding protein. $wt^{Sd}$ ($Sd^{223-440}$), $Asp276Ala^{Sd}$, $Tyr435His^{Sd}$ and $wt^{TEAD4}$ ($TEAD4^{217-434}$) were purified as mixtures of myristoylated (+ Myr) and palmitoylated (+ Palm) proteins.

**Figure S4.** Circular dichroism analysis of $wt^{Sd}$, $Asp276Ala^{Sd}$ and $Tyr435His^{Sd}$. The proteins were dialysed in 20 mM phosphate buffer pH 7.4, 100 mM KF, 0.25 mM TCEP and diluted in this buffer to 0.2 $mg.ml^{-1}$ and far-UV CD spectra were recorded as previously described (Mesrouze et al., 2017a). **A.** CD spectra of $TEAD^{217-434}$ (red) and $wt^{Sd}$ (blue). **B.** CD spectra of $wt^{Sd}$ (blue), $Asp276Ala^{Sd}$ (red) and $Tyr435His^{Sd}$ (green). The represented spectra are the average of two separate measurements.

**Figure S5.** Surface Plasmon Resonance. In all the experiments, N-biotinylated-Avitagged $wt^{Sd}$ was immobilized on sensor chips. **A-L**. For the $K_d$ determinations made at equilibrium, the experiments were conducted as previously described (Mesrouze et al., 2017a). The flow rate was decreased to 20 µl/min with $Vg^{288-337}$, $VGLL2^{85-108}$ and Flyman to get a longer association time. The data were fitted with the Biacore T200 evaluation software using a 1:1 binding model with background. The upper and lower panels show representative sensorgrams and the corresponding binding isotherms. **M-Q**. For the kinetic titration experiments, five analyte concentrations were injected sequentially (100 to 400 s) with a short dissociation between each injection. The last injection was followed by a 500 to 12500 s dissociation step. The data were fitted with the Single Cycle Kinetic module of the Biacore T200 evaluation software using a 1:1 binding model. For all the figures, the signal measured at equilibrium ($R_{max}^{eq}$) and the maximal feasible signal ($R_{max}^{th}$) are given.

**Figure S6**. Superimposition of the structure of the $Vg^{298-337}$:$Sd^{223-440}$ complex on that of the YAP:TEAD complex. $Sd^{223-440}$ (PDB 6Y20) and $TEAD4^{217-434}$ (PDB 6GE3 (Mesrouze et al., 2018)) are represented by green and magenta ribbons, respectively. Dots represent regions that could not be solved. The fatty acid present in the central cavity of Sd or TEAD4 is represented by sticks. The fatty acid was modelled as covalently bound to $Lys350^{Sd}$ or non-covalently bound to TEAD4. The figure was drawn with PyMOL (Schrödinger Inc., Cambridge, MA).

**Figure S7**. Primary sequence alignments of Vg proteins from different insect species. The primary sequences were manually aligned using the sequence of Vg from *D. melanogaster* as reference. The secondary structure adopted by Vg upon binding to Sd is indicated. α: α-helix; **Ω**: **Ω**-loop. Gaps and conserved residues are indicated by dashes and asterisks, respectively. *D. melanogaster* NP_523723; *G. pallidipes* A0A1B0A9G4; *B. olea* XP_014095034; *C. marinus* A0A1J1IJG6; *A. mellifera* XP_001122002; *A. colombica* XP_018046781; *C. cinctus* XP_015601999; *M. demolitor* XP_008553424; *P. machaon* XP_014359924; *H. armigera* XP_021192233; *S. litura* XP_022834734; *H. virescens* A0A2A4K2F4; *P. pyralis* A0A1Y1MB82; *O. taurus* XP_022913656; *T. castaneum* XP_008199328; *A. tumida* XP_019864741.

**Figure S8**. TR-FRET assay. The figure shows representative inhibition curves obtained in the TR-FRET assay. Biotinylated N-Avitagged-$TEAD4^{217-434}$ (1 nM) and LANCE Eu-W1024

Streptavidin (0.5nM, PerkinElmer, Waltham, MA) were pre-incubated for 1 h at room temperature in 50 mM HEPES pH 7.4, 100 mM KCl, 0.05% (v/v) Tween-20, 0.25 mM TCEP, 1 mM EDTA, and 0.05% (w/v) BSA. After incubation, N-terminally Cy5-labelled $YAP^{60-100}$ (20 nM) and serial dilutions of the peptides/protein fragments to be tested were added and incubated in white 384-well plates (Greiner Bio-One International, Austria) for 1 h at room temperature. DMSO or acetonitrile was present at 2% in the assay. The fluorescence was measured with a Genios Pro reader (Tecan, Switzerland) (50 µs delay between excitation and fluorescence, 75 µs integration time, excitation wavelength of 340 nm, emission wavelengths of 620 nm and 665 nm). Twelve stepwise dilutions (dilution factor 3.33) of each peptide were used in the experiments. The highest concentrations present in the assay were 222 µM for $Vg^{323-337}$, $VGLL2^{135-149}$ and $VGLL3^{140-154}$ or 250 µM for $YAP^{85-99}$. The solubility of the peptides in the assay buffer was measured with a NEPHELOStar (BMG Labtech GmbH., Germany). Data analyses were carried out using the TR-FRET 655/620 nm emission ratio. The $IC_{50}$ values were estimated by fitting the data by nonlinear fit regression with GraphPad Prism (GraphPad Software, San Diego, CA). The asterisks correspond to 100% inhibition. No $IC_{50}$ value could be determined for $VGLL3^{140-154}$. The $IC_{50}$ curves for $Vg^{288-337}$ and Flyman are not represented because they were lower than the TEAD4 concentration present in the assay.

**Figure S1**

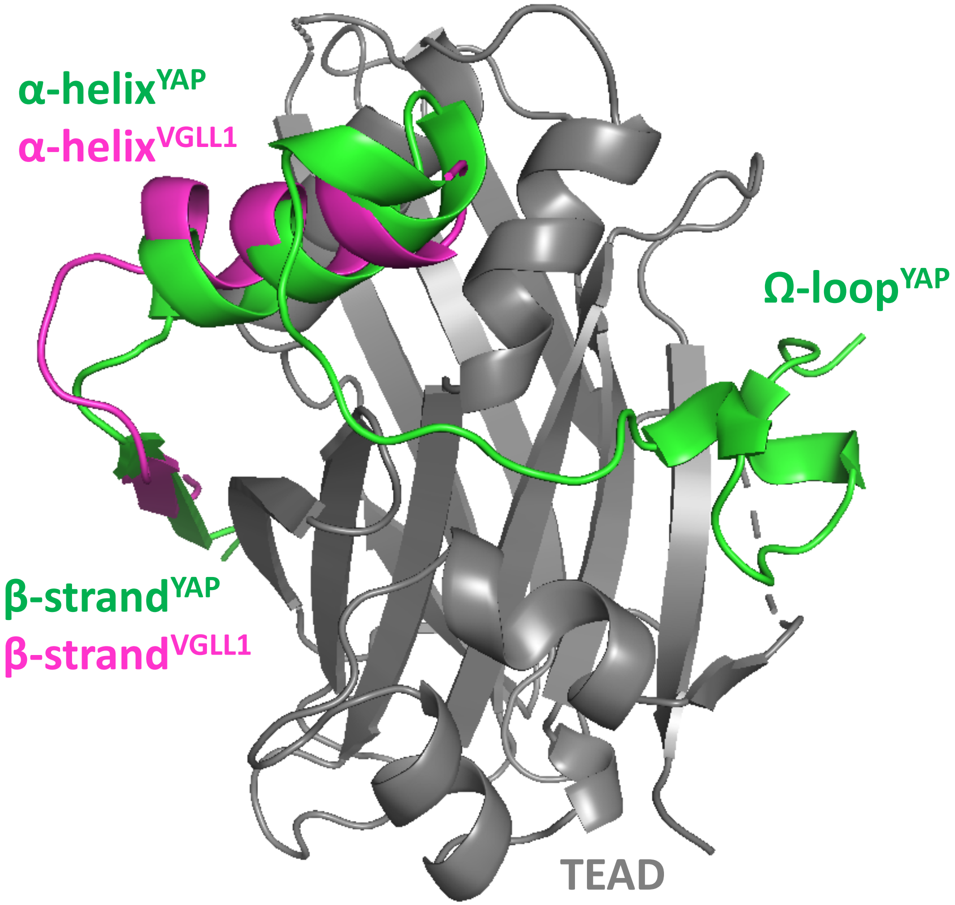
α-helix$^{YAP}$
α-helix$^{VGLL1}$
Ω-loop$^{YAP}$
β-strand$^{YAP}$
β-strand$^{VGLL1}$
TEAD

### Figure S2

```
Sd223-440        RAIATHKFRLLEFTAFMEIQRD-EIYHRHLFVQLGG-KPSFSDPLLETVD
TEAD4217-434     RSVASSKLWMLEFSAFLEQQQDPDTYNKHLFVHIGQSSPSYSDPYLEAVD
                 *::*: *: :***:**:* *:* : *::****::*  .**:*** **:**
Sd223-440        IRQIFDKFPEKSGGLKDLYEKGPQNAFYLVKCWADLNTDLTTGSETGDFY
TEAD4217-434     IRQIYDKFPEKKGGLKDLFERGPSNAFFLVKFWADLNTNIE--DEGSSFY
                 ****:******.******:*:**.***:*** ******::   .* ..**
Sd223-440        GVTSQYESNENVVLVCSTIVCSFGKQVVEKVESEYSRLENNRYVYRIQRS
TEAD4217-434     GVSSQYESPENMIITCSTKVCSFGKQVVEKVETEYARYENGHYSYRIHRS
                 **:***** **:::.*** *************:**:* **.:* ***:**
Sd223-440        PMCEYMINFIQKLKNLPERYMMNSVLENFTILQVMRARETQETLLCIAYV
TEAD4217-434     PLCEYMINFIHKLKHLPEKYMMNSVLENFTILQVVTNRDTQETLLCIAYV
                 *:********:***:***:***************:  *:***********
Sd223-440        FEVAAQNSGTTHHIYRLIKE
TEAD4217-434     FEVSASEHGAQHHIYRLVKE
                 ***:*.: *: ******:**
```

#### Figure S3

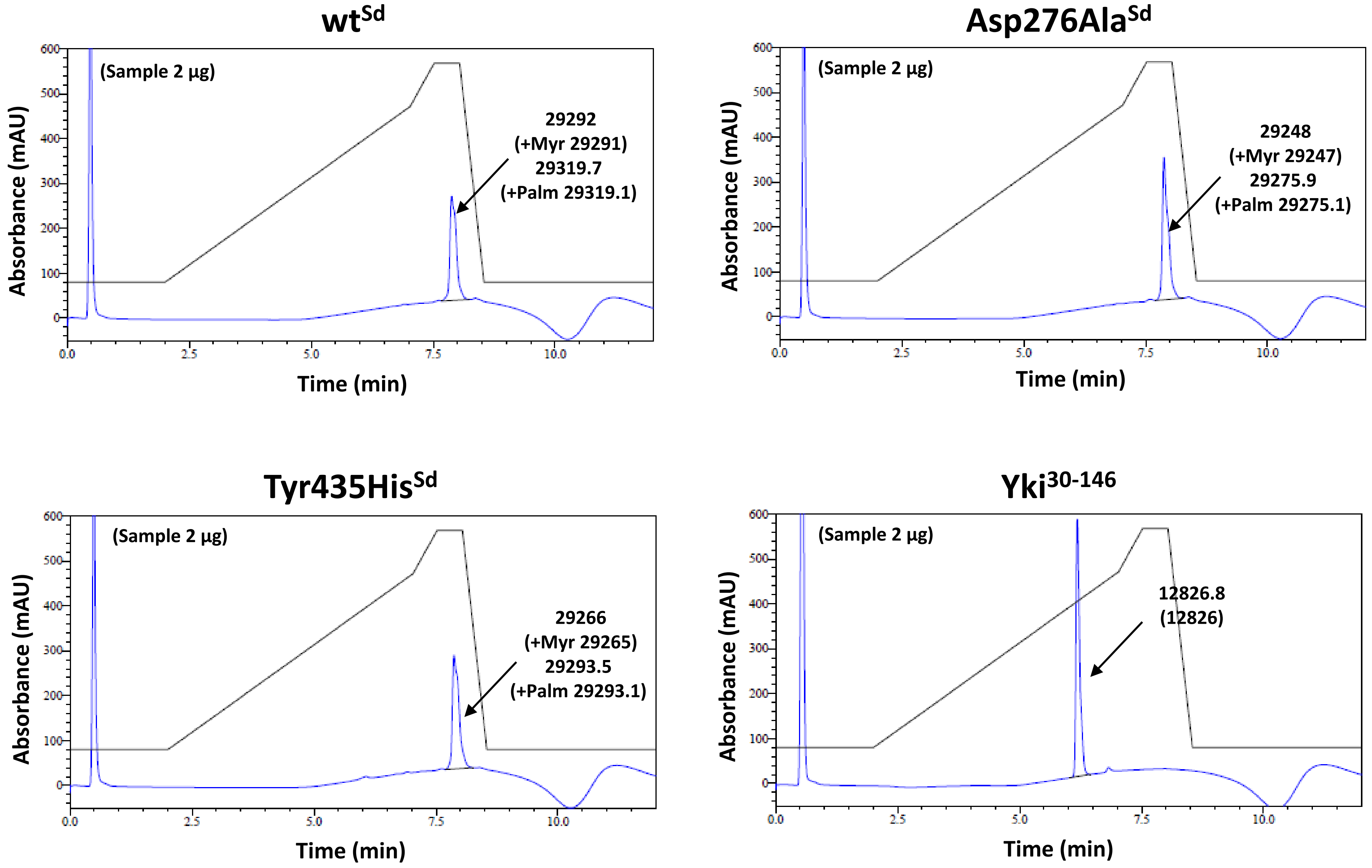

wtSd
(Sample 2 µg)
29292
(+Myr 29291)
29319.7
(+Palm 29319.1)
Absorbance (mAU)
Time (min)
Asp276AlaSd
(Sample 2 µg)
29248
(+Myr 29247)
29275.9
(+Palm 29275.1)
Absorbance (mAU)
Time (min)
Tyr435HisSd
(Sample 2 µg)
29266
(+Myr 29265)
29293.5
(+Palm 29293.1)
Absorbance (mAU)
Time (min)
Yki30-146
(Sample 2 µg)
12826.8
(12826)
Absorbance (mAU)
Time (min)

### Figure S3

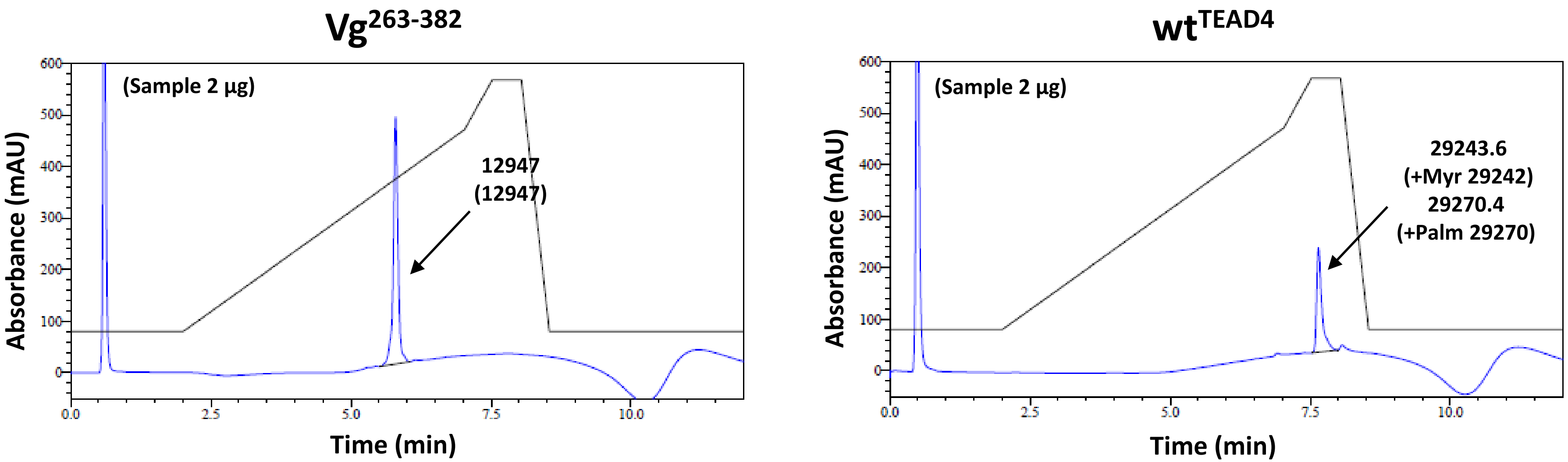
Vg263-382
(Sample 2 µg)
12947
(12947)
Absorbance (mAU)
Time (min)
wtTEAD4
(Sample 2 µg)
29243.6
(+Myr 29242)
29270.4
(+Palm 29270)
Absorbance (mAU)
Time (min)

#### Figure S4

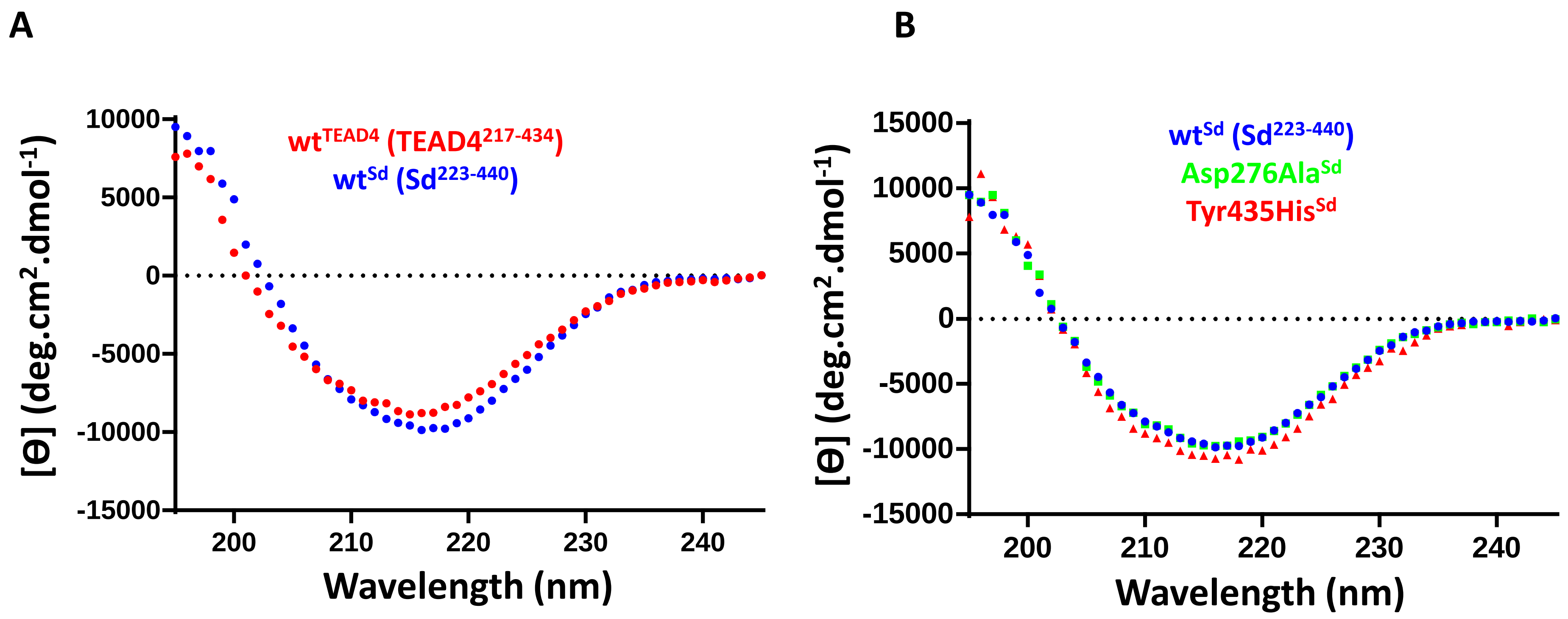

A
wt$^{TEAD4}$ (TEAD4$^{217-434}$)
wt$^{Sd}$ (Sd$^{223-440}$)
[Θ] (deg.cm$^2$.dmol$^{-1}$)
10000
5000
0
-5000
-10000
-15000
200
210
220
230
240
Wavelength (nm)
B
wt$^{Sd}$ (Sd$^{223-440}$)
Asp276Ala$^{Sd}$
Tyr435His$^{Sd}$
[Θ] (deg.cm$^2$.dmol$^{-1}$)
15000
10000
5000
0
-5000
-10000
-15000
200
210
220
230
240
Wavelength (nm)

### Figure S5

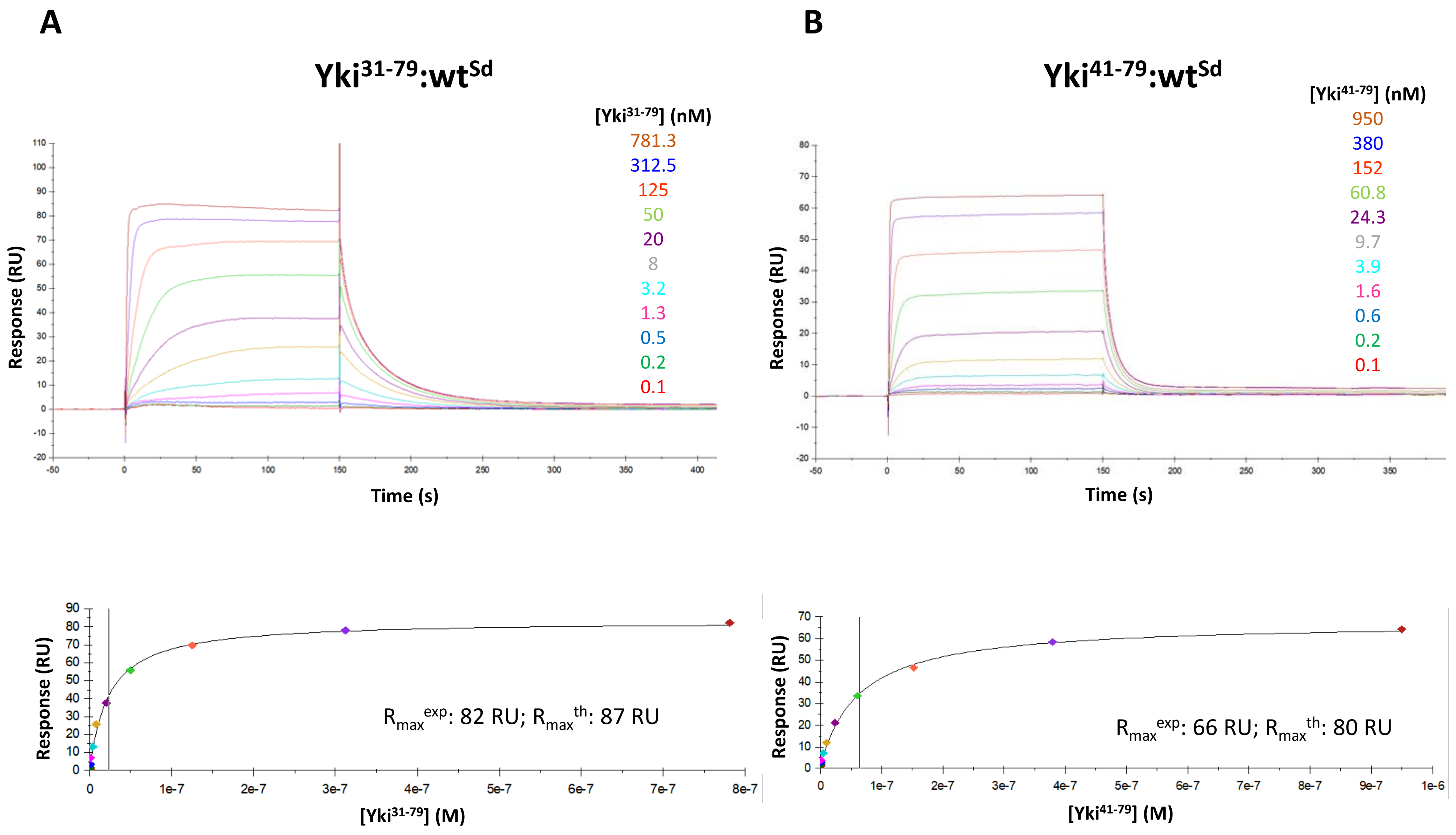
A
Yki31-79:wtSd
[Yki31-79] (nM)
781.3
312.5
125
50
20
8
3.2
1.3
0.5
0.2
0.1
Response (RU)
Time (s)
Rmax exp: 82 RU; Rmax th: 87 RU
[Yki31-79] (M)
B
Yki41-79:wtSd
[Yki41-79] (nM)
950
380
152
60.8
24.3
9.7
3.9
1.6
0.6
0.2
0.1
Response (RU)
Time (s)
Rmax exp: 66 RU; Rmax th: 80 RU
[Yki41-79] (M)

#### Figure S5

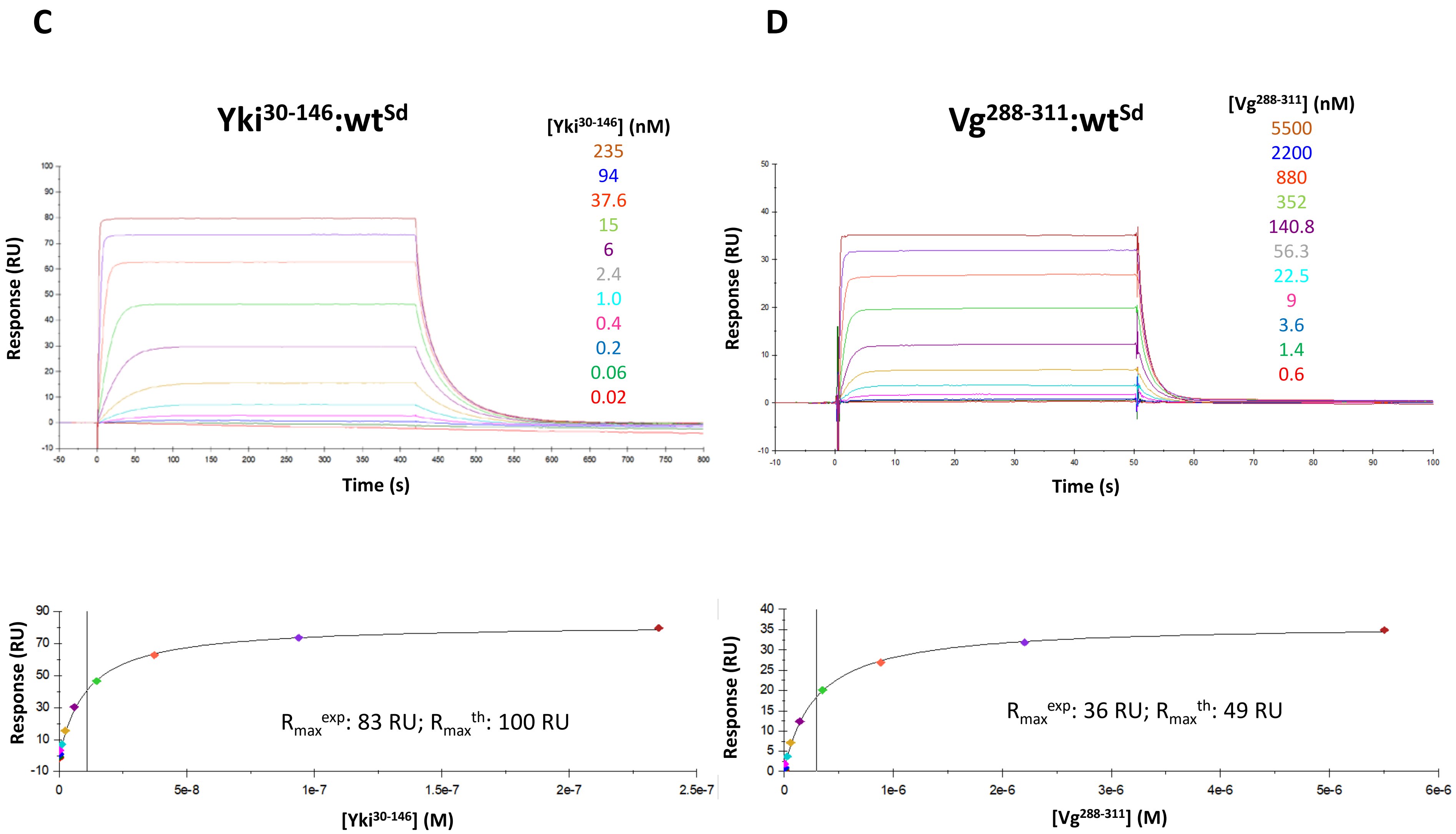
C
D
Yki30-146:wtSd
[Yki30-146] (nM)
235
94
37.6
15
6
2.4
1.0
0.4
0.2
0.06
0.02
Response (RU)
Time (s)
Vg288-311:wtSd
[Vg288-311] (nM)
5500
2200
880
352
140.8
56.3
22.5
9
3.6
1.4
0.6
Rmaxexp: 83 RU; Rmaxth: 100 RU
[Yki30-146] (M)
Rmaxexp: 36 RU; Rmaxth: 49 RU
[Vg288-311] (M)

### Figure S5

E

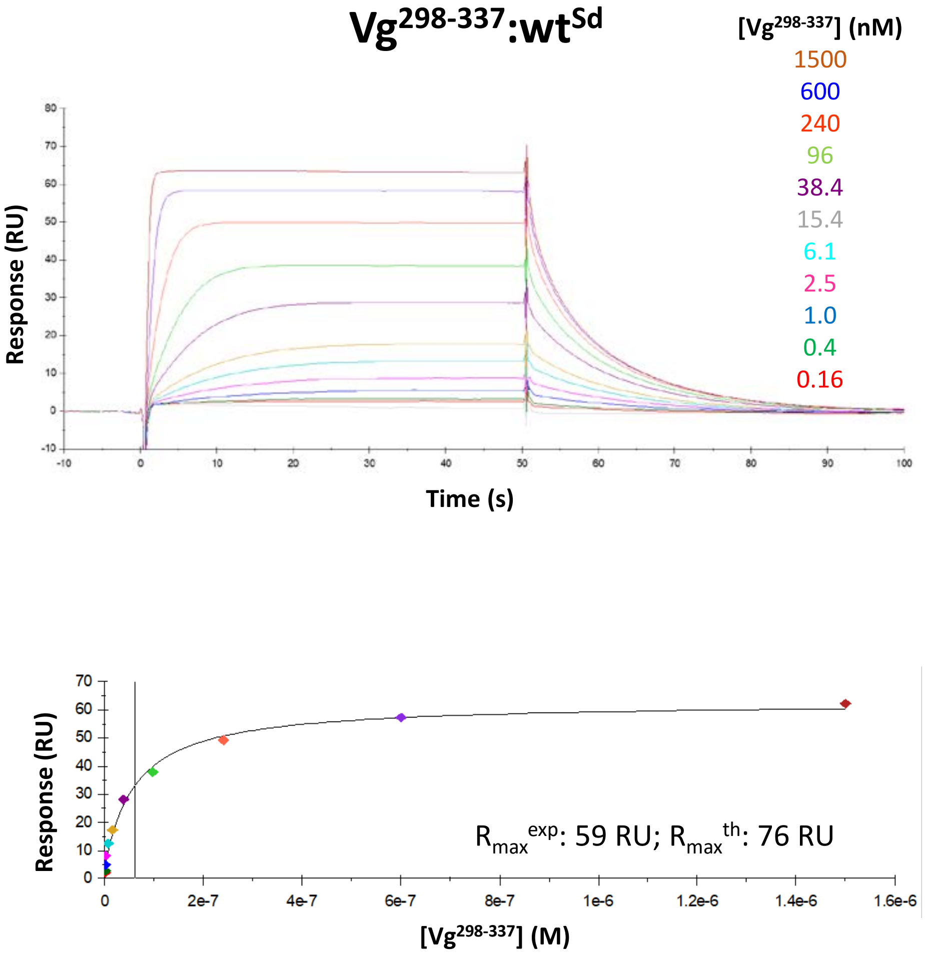
Vg298-337:wtSd
[Vg298-337] (nM)
1500
600
240
96
38.4
15.4
6.1
2.5
1.0
0.4
0.16
Response (RU)
Time (s)
Rmax exp: 59 RU; Rmax th: 76 RU
[Vg298-337] (M)

F

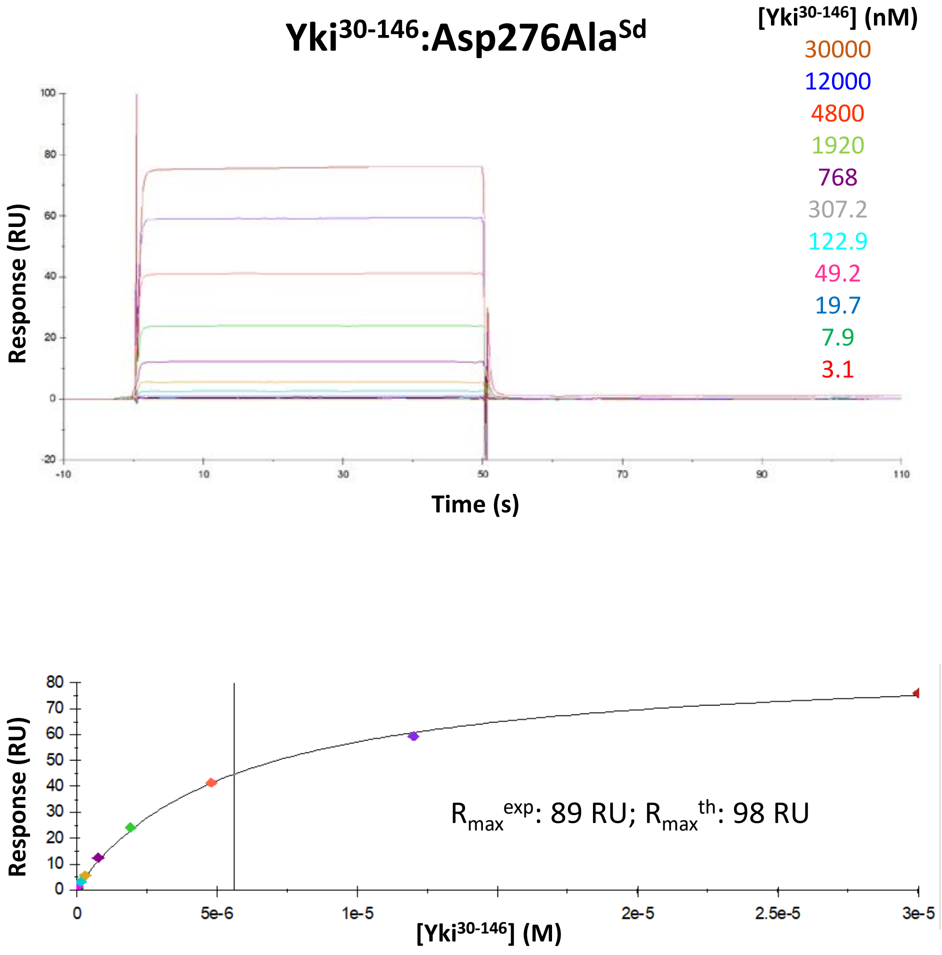
Yki30-146:Asp276AlaSd
[Yki30-146] (nM)
30000
12000
4800
1920
768
307.2
122.9
49.2
19.7
7.9
3.1
Response (RU)
Time (s)
Rmax exp: 89 RU; Rmax th: 98 RU
[Yki30-146] (M)

### Figure S5

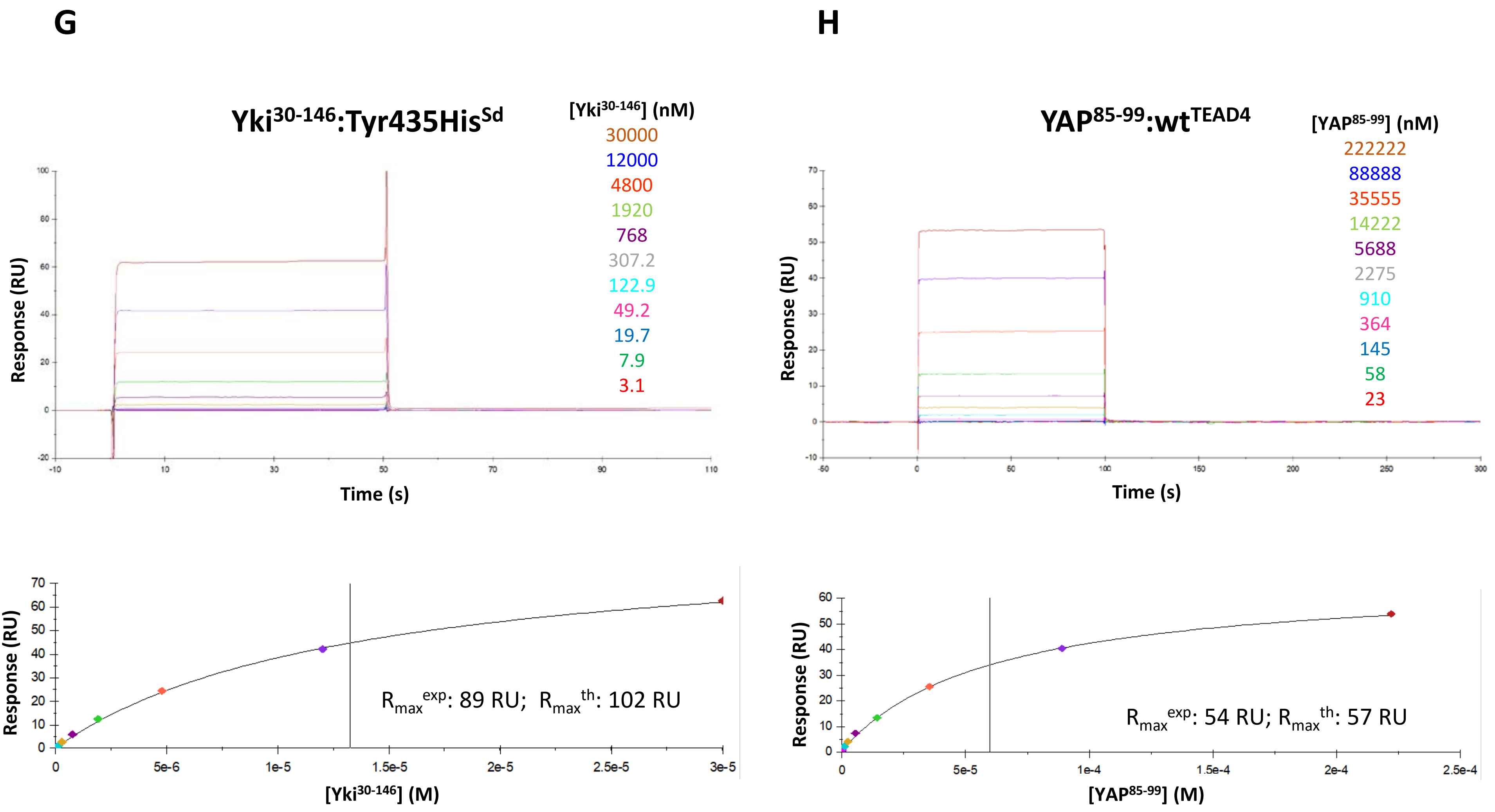

G
Yki30-146:Tyr435HisSd
[Yki30-146] (nM)
30000
12000
4800
1920
768
307.2
122.9
49.2
19.7
7.9
3.1
Response (RU)
Time (s)
Rmaxexp: 89 RU; Rmaxth: 102 RU
[Yki30-146] (M)
H
YAP85-99:wtTEAD4
[YAP85-99] (nM)
222222
88888
35555
14222
5688
2275
910
364
145
58
23
Response (RU)
Time (s)
Rmaxexp: 54 RU; Rmaxth: 57 RU
[YAP85-99] (M)

### Figure S5

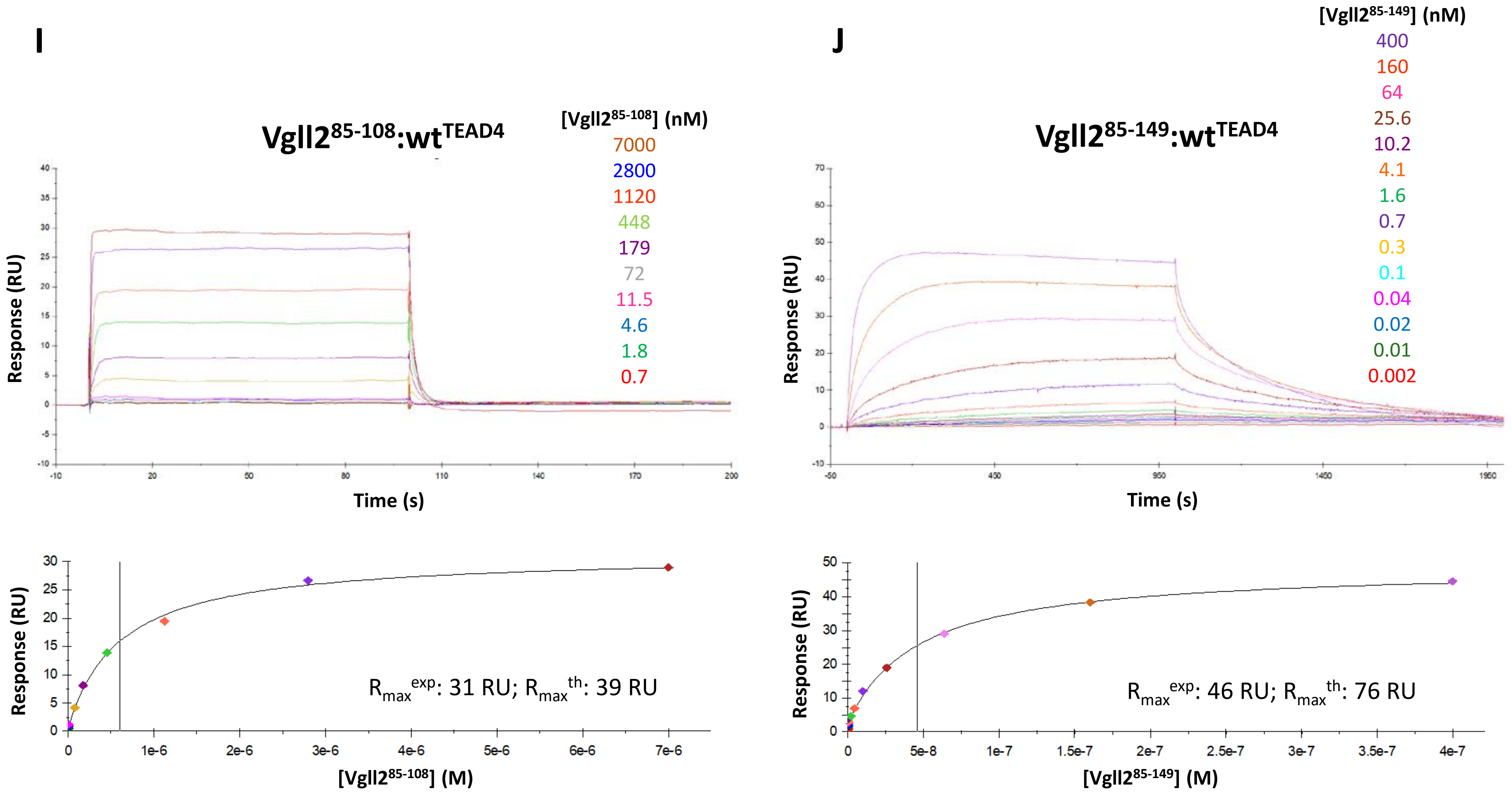
I
Vgll2^85-108:wt^TEAD4
[Vgll2^85-108] (nM)
7000
2800
1120
448
179
72
11.5
4.6
1.8
0.7
Response (RU)
Time (s)
R_max^exp: 31 RU; R_max^th: 39 RU
[Vgll2^85-108] (M)
J
Vgll2^85-149:wt^TEAD4
[Vgll2^85-149] (nM)
400
160
64
25.6
10.2
4.1
1.6
0.7
0.3
0.1
0.04
0.02
0.01
0.002
Response (RU)
Time (s)
R_max^exp: 46 RU; R_max^th: 76 RU
[Vgll2^85-149] (M)

### Figure S5

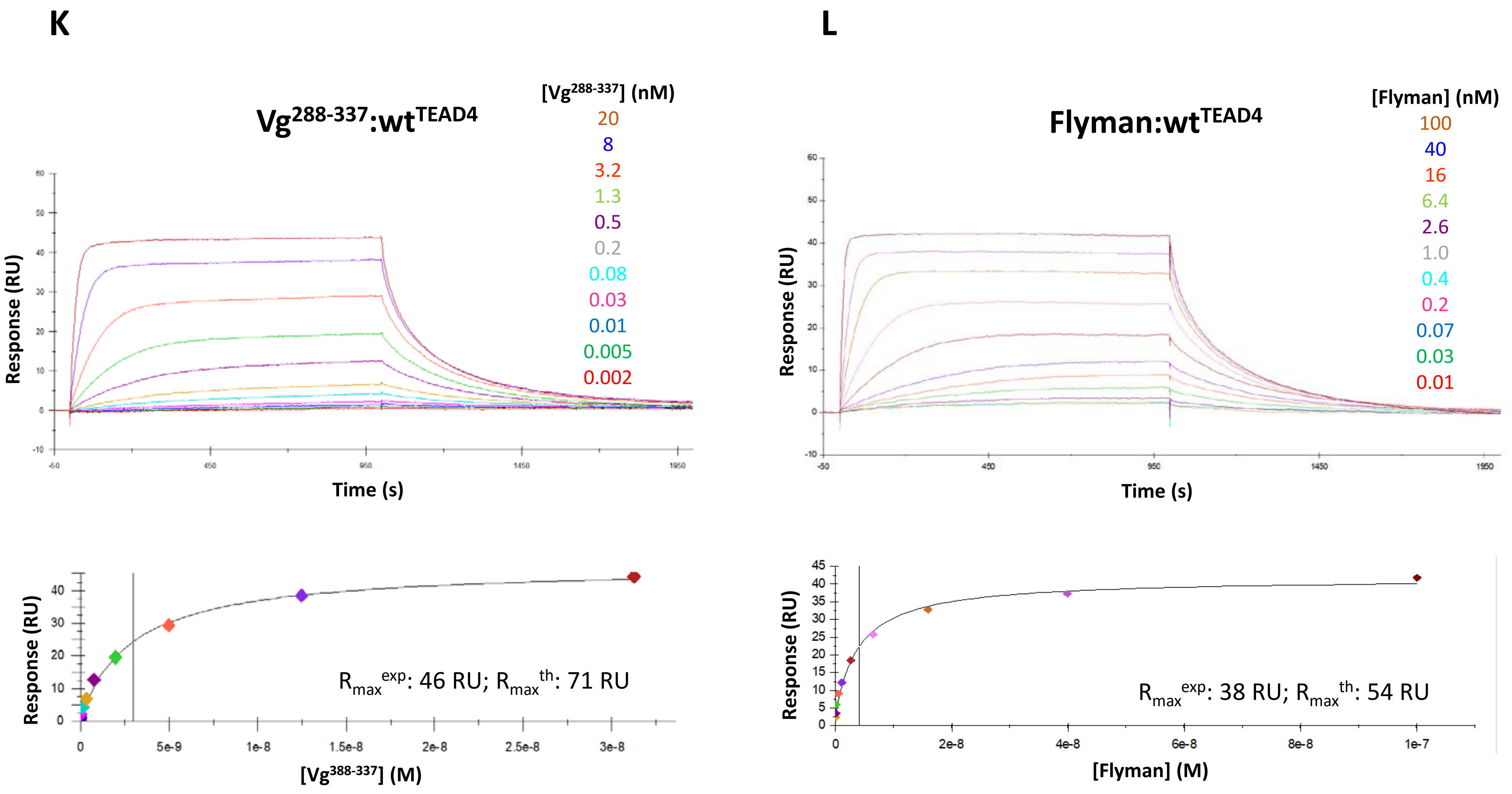
K
Vg288-337:wtTEAD4
[Vg288-337] (nM)
20
8
3.2
1.3
0.5
0.2
0.08
0.03
0.01
0.005
0.002
Response (RU)
Time (s)
Rmax exp: 46 RU; Rmax th: 71 RU
[Vg388-337] (M)
L
Flyman:wtTEAD4
[Flyman] (nM)
100
40
16
6.4
2.6
1.0
0.4
0.2
0.07
0.03
0.01
Response (RU)
Time (s)
Rmax exp: 38 RU; Rmax th: 54 RU
[Flyman] (M)

### Figure S5

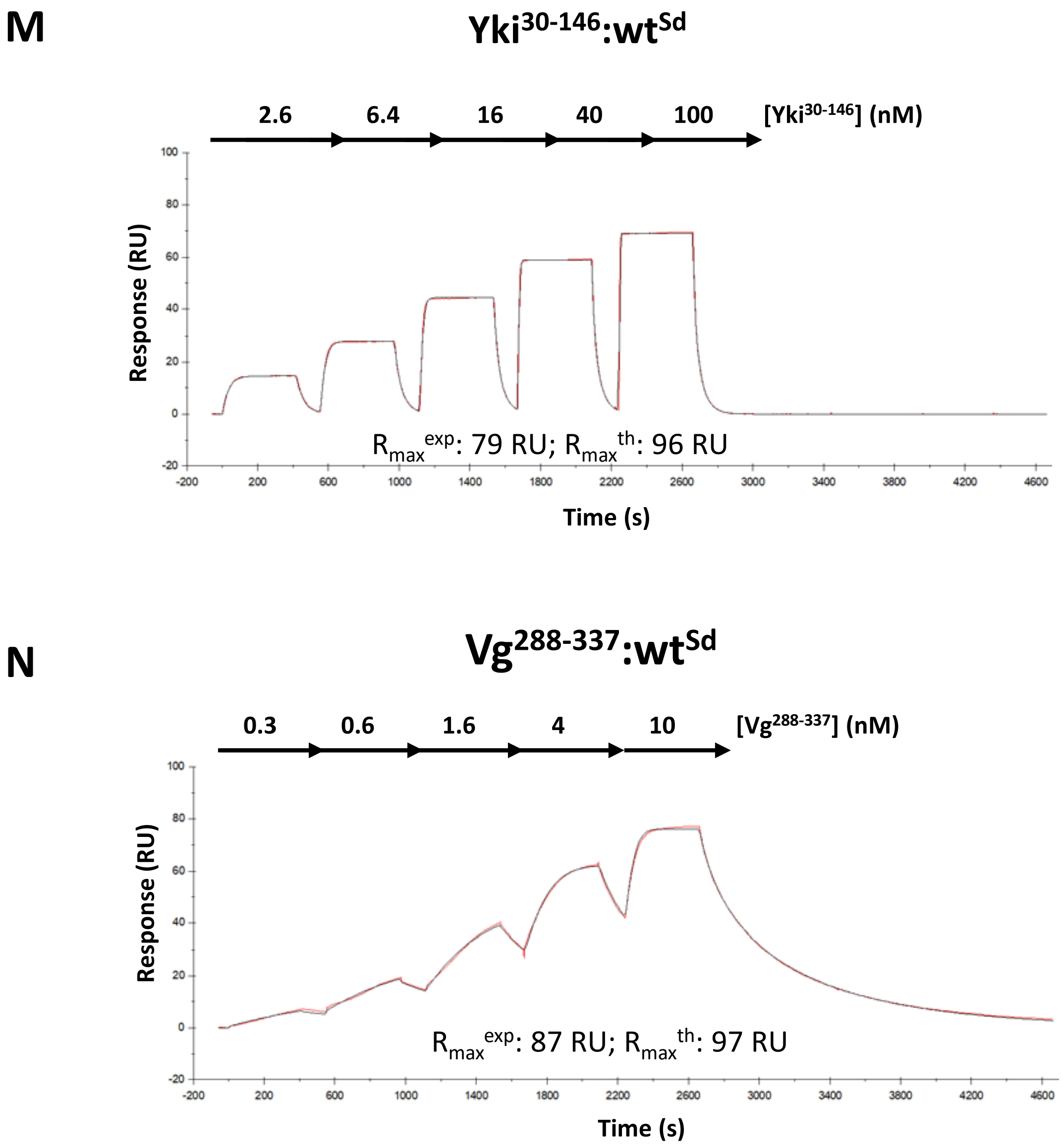
M
$Yki^{30-146}$:$wt^{Sd}$
2.6
6.4
16
40
100
[$Yki^{30-146}$] (nM)
Response (RU)
$R_{max}^{exp}$: 79 RU; $R_{max}^{th}$: 96 RU
Time (s)
N
$Vg^{288-337}$:$wt^{Sd}$
0.3
0.6
1.6
4
10
[$Vg^{288-337}$] (nM)
Response (RU)
$R_{max}^{exp}$: 87 RU; $R_{max}^{th}$: 97 RU
Time (s)

#### Figure S5

**O** **$Vg^{263\text{-}382}$:$wt^{Sd}$**

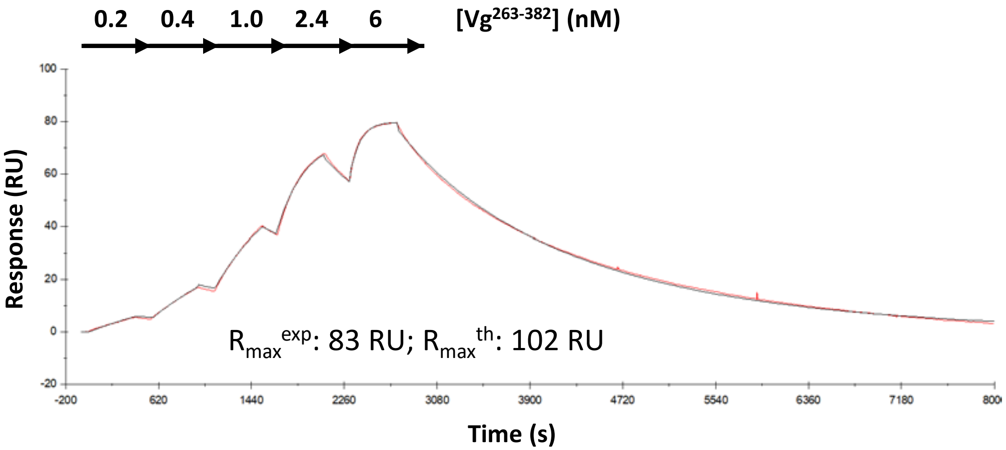


**P** **$Vg^{263\text{-}382}$:$Asp276Ala^{Sd}$**

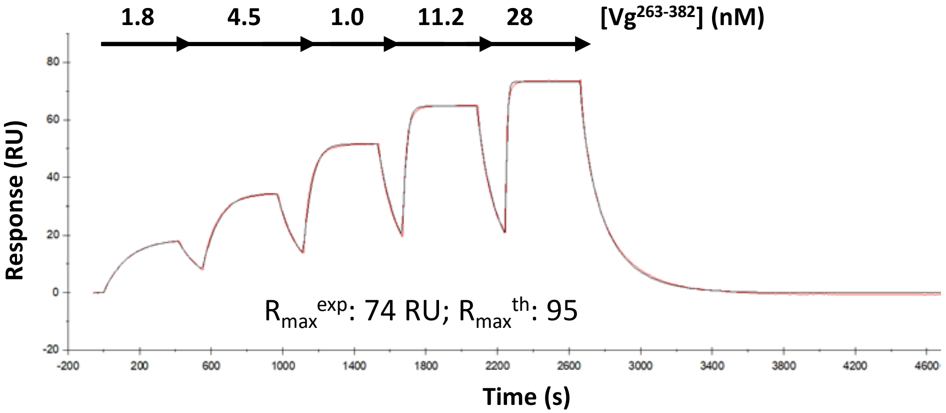

### Figure S5

**Q**

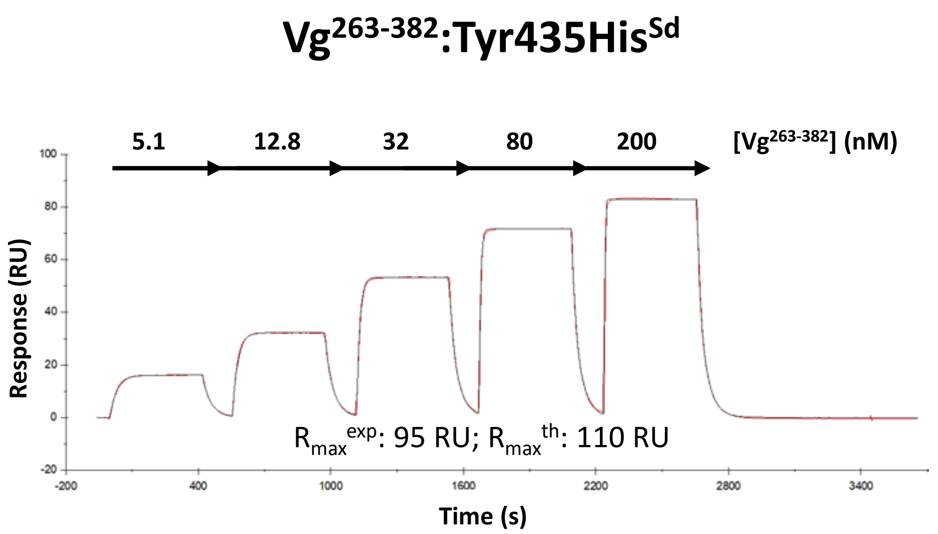

Vg263-382:Tyr435HisSd
5.1
12.8
32
80
200
[Vg263-382] (nM)
Response (RU)
100
80
60
40
20
0
-20
-200
400
1000
1600
2200
2800
3400
Time (s)
Rmaxexp: 95 RU; Rmaxth: 110 RU

**Figure S6**

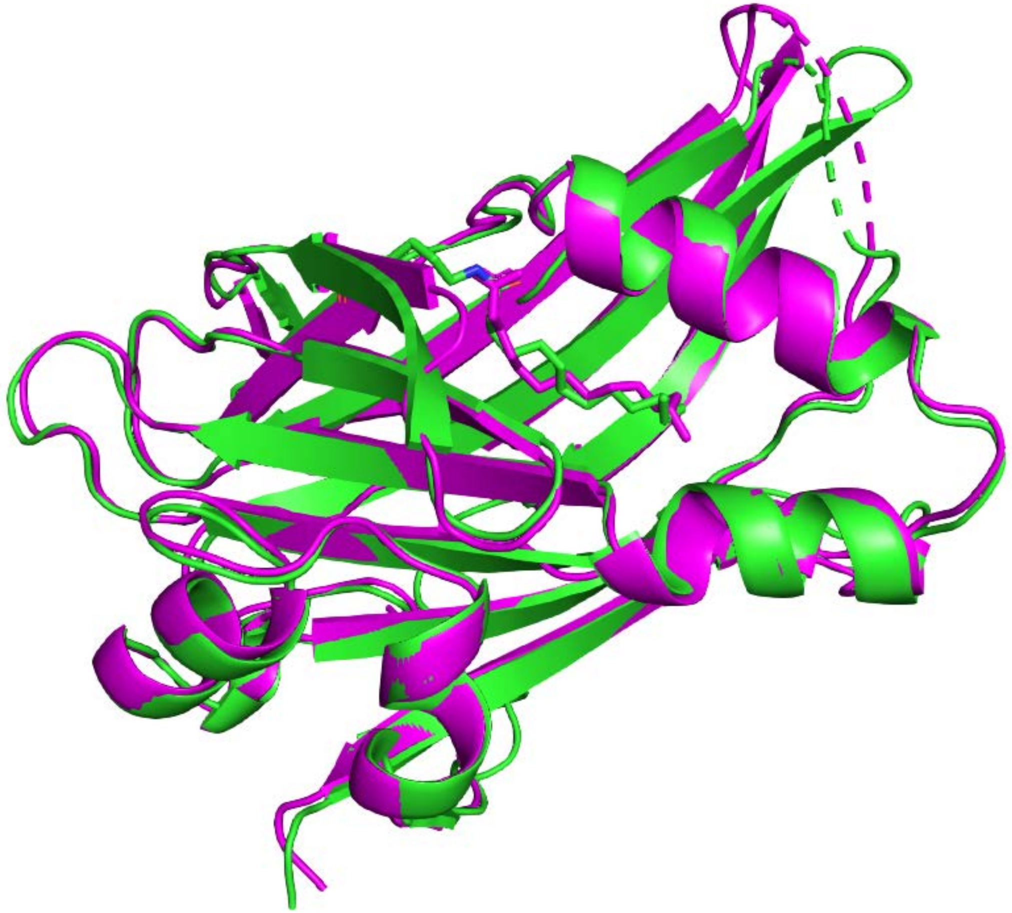

**Figure S7**

```
                              ααααααααααααα            ΩΩΩΩΩΩΩΩΩΩΩΩΩΩΩ
Diptera
Drosophila melanogaster       CVVFTNYSGDTASQVDEHFSRALNYNNKDSKESSSPMSNRNFPPSFWNSN
Glossina pallidipes           CVVFTNYSGDTASVVDEHFSRALNFNNKDSKEASSPMSARNFPPSFWNSN
Bactrocera oleae              CVVFTNYSGDTASVVDEHFSRALNFNNKDSKETGSPMSARNFPPSFWNSN
Clunio marinus                CVVVTHYSGDVANVVDEHFTRALNFNDKNNKDSN-PMSARNFPPSFWNSN
Hymenoptera
Apis mellifera                CVVFTHYRGDAASEVEEHFQRALAHDKPKENIS--PMFMRNFPPSFWKSQ
Microplitis demolitor         CVVFTHYRGDAASEVEEHFQRALAHDKVKENI---PMSTRNFPPSFWNSQ
Atta colombica                CVVFTHYRGDAASEVEEHFQRALAHDKLKENIS--PMSMRNFPPSFWNSQ
Cephus cinctus                CVVFTHYRGDAATEVEEHFQRALAHDKLKENIS--PMAMRNFPPSFWNSQ
Lepidoptera
Papilio machaon               CVVFTHYAGDVAAVVDEHFARALAIDKPKESV---PLASRNLPASFFNAA
Helicoverpa armigera          CVVFTHYSGDVAAVVDEHFARALSLDKPKESV---PMVSRNLPASFFNAA
Spodoptera litura             CVVFTHYSGDVAAVVDEHFARALSLDKPKESV---PMVSRNLPASFFNAA
Heliothis virescens           CVVFTHYSGDVAAVVDEHFARALSLDKPKESV---PMVSRNLPASFFNAA
Coleoptera
Photinus pyralis              CVVFTHYQGDASSVVDEHFSRALDKTSQTKESP--PMSQRNFPPSFWNCQ
Onthophagus taurus            CVVFTHYQGDTASVVDEHFSRALDKNNHAKEIS--PMSTRNFPASFWNSQ
Tribolium castaneum           CVVFTHYQGDAASVVDEHFSRALDKTSTHTKESS-PMSARNFPPSFWDSQ
Aethina tumida                CVVFTHYQGDAATVVDEHFSRALDKTAAHTKESS-PMSSRNFPPSFWNSQ
                              *** * * **    * *** ***                ** * **
```

Figure S8

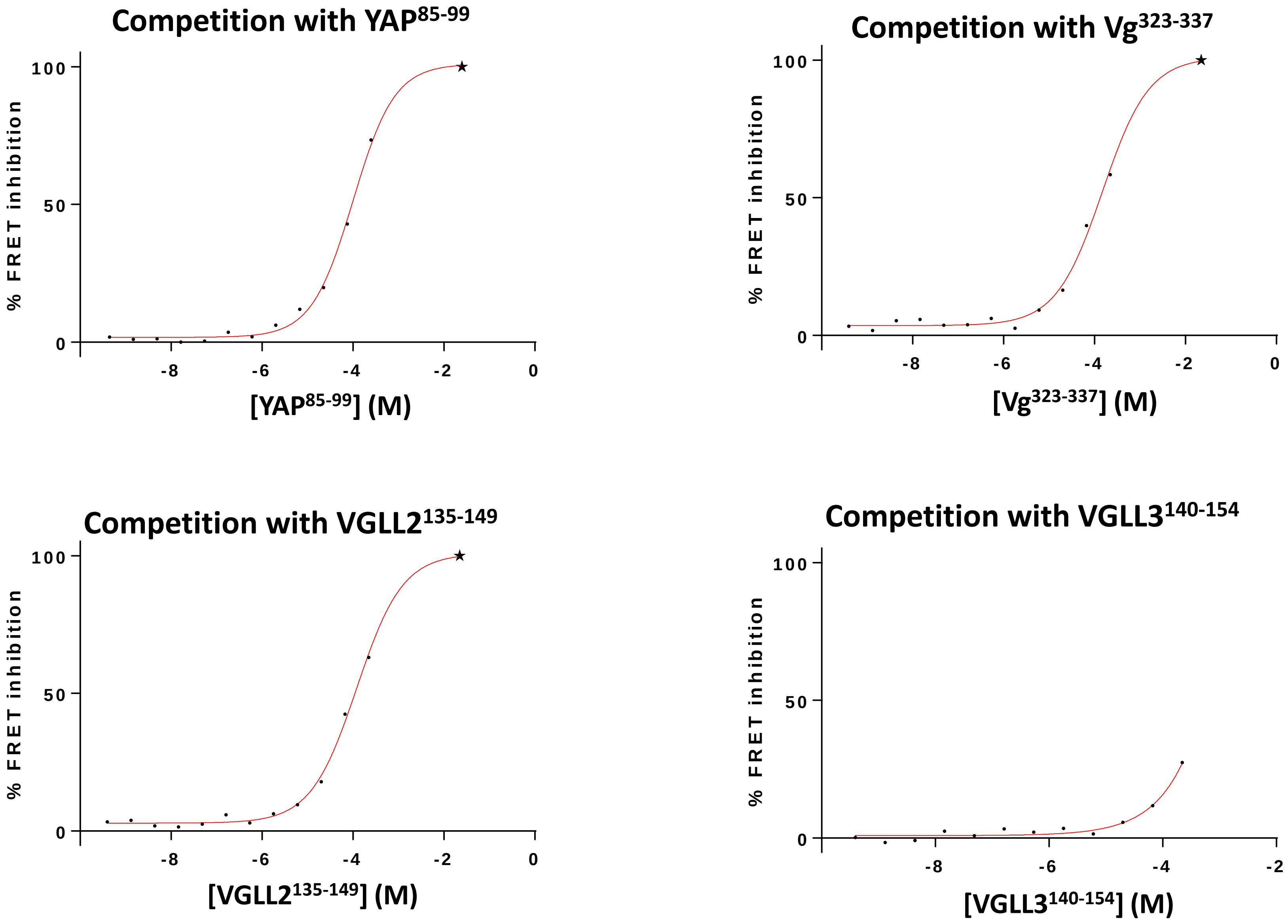

Competition with YAP85-99
100
50
0
% FRET inhibition
-8
-6
-4
-2
0
[YAP85-99] (M)
Competition with Vg323-337
100
50
0
% FRET inhibition
-8
-6
-4
-2
0
[Vg323-337] (M)
Competition with VGLL2135-149
100
50
0
% FRET inhibition
-8
-6
-4
-2
0
[VGLL2135-149] (M)
Competition with VGLL3140-154
100
50
0
% FRET inhibition
-8
-6
-4
-2
[VGLL3140-154] (M)

### Figure S8

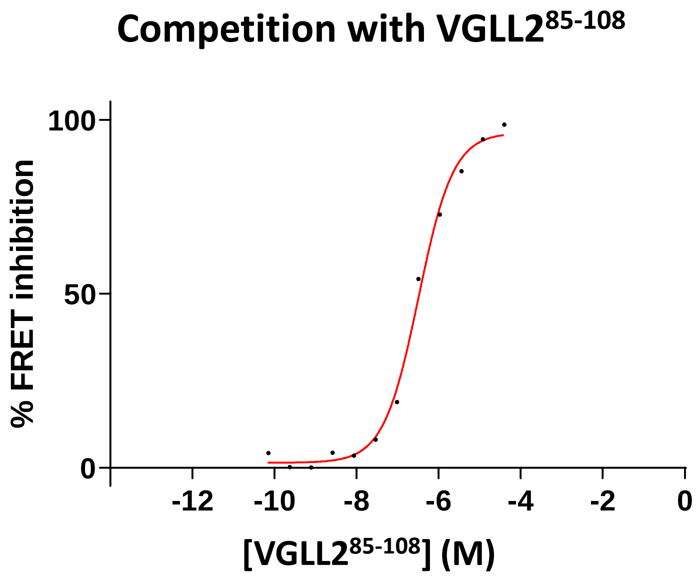
Competition with VGLL2$^{85-108}$
% FRET inhibition
100
50
0
-12
-10
-8
-6
-4
-2
0
[VGLL2$^{85-108}$] (M)

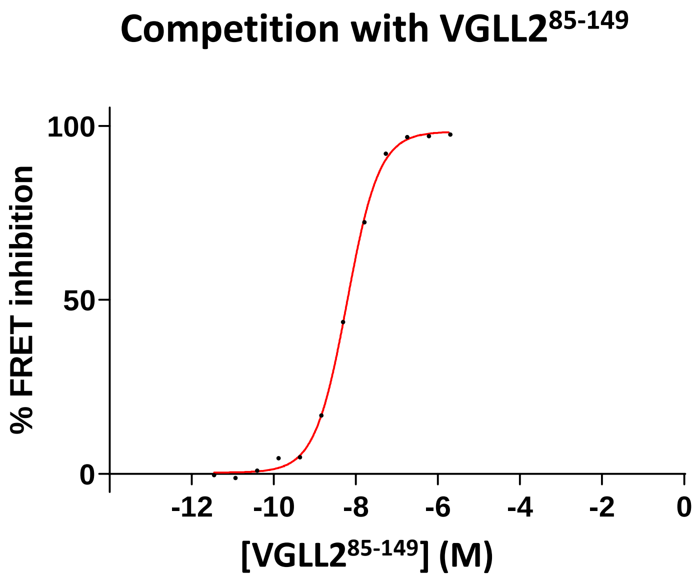
Competition with VGLL2$^{85-149}$
% FRET inhibition
100
50
0
-12
-10
-8
-6
-4
-2
0
[VGLL2$^{85-149}$] (M)
